# Targeted memory reactivation during post-learning sleep does not enhance motor memory consolidation in older adults

**DOI:** 10.1101/2022.10.13.512106

**Authors:** Judith Nicolas, Julie Carrier, Stephan P. Swinnen, Julien Doyon, Geneviève Albouy, Bradley R. King

## Abstract

Targeted memory reactivation (TMR) during sleep enhances memory consolidation in young adults by modulating electrophysiological markers of neuroplasticity. Interestingly, older adults exhibit deficits in motor memory consolidation, an impairment that has been linked to age-related degradations in the same sleep features sensitive to TMR. We hypothesized that TMR would enhance consolidation in older adults via the modulation of these markers. Seventeen older participants were trained on a motor task involving two auditory-cued sequences. During a post-learning nap, two auditory cues were played: one associated to a learned (i.e., reactivated) sequence and one control. Performance during two delayed retests did not differ between reactivated and non-reactivated sequences. Moreover, both associated and control sounds modulated brain responses, yet there were no consistent differences between the auditory cue types. Our results collectively demonstrate that older adults do not benefit from specific reactivation of a motor memory trace by an associated auditory cue during post-learning sleep. It is possible, however, that auditory stimulation during post-learning sleep boosts motor memory consolidation in a non-specific manner.

## 1. Introduction

The development and maintenance of the incredibly vast repertoire of human movement depends on motor plasticity processes, which facilitates the learning, consolidation, and retrieval of motor skills. There is considerable evidence in young adults indicating that sleep following initial learning provides a privileged window for the newly acquired memory to be consolidated into a stable, longer-term trace [1, 2]. Similar sleep-related benefits appear to be predominantly absent in healthy older adults [3, 4, 5, 6 but see 7, 8], a result that is, at least partially, linked to age-related degradations in sleep macro- and micro-architecture [5, 9]. Accordingly, the development and implementation of interventions enhancing sleep-facilitated memory consolidation in older adults offers a promising avenue to alleviate aging-associated deficits in learning and memory processes.

One approach that has the potential to enhance sleep-facilitated consolidation is targeted memory reactivation (TMR) [10, 11, 11]. Briefly, TMR consists of associating a sensory stimulus (e.g., a sound) with to-be-learned material (e.g., word pairs or a sequence of movements) during an initial encoding session. This stimulus is subsequently replayed during a post-learning sleep episode and is thought to reactivate the recently acquired memory trace. In young adults, TMR has proven to be effective in boosting consolidation processes following declarative [e.g. 13, 14, 15, 16] and motor [e.g. 17, 18, 19, 20] learning. These beneficial effects at the behavioral level are paralleled by specific modulations of electrophysiological markers of plasticity during sleep. Specifically, our recent research in the motor memory domain in young adults revealed that TMR increases Slow Wave (SW – high amplitude waves in the 0.5–2 Hz frequency band) characteristics, such as density and peak-to-peak (PTP) amplitude. Additionally, an increase in coupling between slow oscillation phase and amplitude of sigma – 12 to 16 Hz – oscillations at the SW peak was related to higher effect of TMR on motor performance. Conversely, sounds that were not associated to learning strengthened SW/sigma coupling during the descending phase of the SW. These results collectively suggested that an increase of SW activity precisely coordinated with sigma oscillations is crucial for motor memory consolidation processes in young adults.

Interestingly, this previous research demonstrated that TMR modulated specific sleep features that have been previously linked to the aforementioned deficits in sleep-related memory consolidation in older adults. For example, aging is known to be accompanied by decreases in NREM 3 sleep duration as well as alterations in sleep-specific electrophysiological markers of plasticity [21]. Indeed, the number of sleep spindles decreases with age, and they exhibit smaller amplitude, shorter duration and lower frequency [5, 22, 23]. Slow wave density and amplitude also decrease [24, 25]; and, the coupling between SW and spindles exhibits age-related degradations [26]. This previous research collectively raises the intriguing possibility that TMR may be an effective avenue to enhance sleep-facilitated consolidation in healthy older adults via the modulation of these sleep features (e.g., slow and sigma oscillations as well as their phase amplitude coupling). The goal of this study therefore was to investigate auditory TMR-induced modulations of motor memory consolidation processes at the behavioral and electrophysiological levels in healthy older adults. We hypothesized that TMR would enhance motor memory consolidation in this population; and this boost would be linked to alterations in slow and sigma oscillation activity, as well as their coupling, during post-learning sleep.

## 2. Materials and Methods

The current study employed nearly identical experimental procedures and data analyses as our previous pre-registered study that examined effects of TMR in healthy young adults [27](only sound presentation devices differed).

### 2.1. Participants

This protocol was approved by the local Ethics Committee (B322201525025) and conducted according to the declaration of Helsinki [28]. Participants gave written informed consent before participating in this study. Monetary compensation was given for participants’ time and effort. Healthy older adults (50 years and older) were recruited from Leuven (Belgium) and the surrounding area to serve as participants. Although our minimum age for being included (50 years) may seem young in the context of aging research, similar cut-offs have been used in previous sleep and motor memory research [3,5,8,29,30]. Moreover, there is evidence that both sleep characteristics [25,31] and sleep-dependent motor memory consolidation[4,29] are significantly altered by 50 years of age. Whereas our analyses in the main text focus on the effects of TMR in older adults at the group level, supplementary Figure S1 includes exploratory analyses assessing age-related changes (between 50 and 74 years of age) within the group of older adults. Additional inclusion criteria were: 1) no previous extensive training with a musical instrument or as a professional typist, 2) free of medical, neurological, psychological, or psychiatric conditions, including depression and anxiety as assessed by the Beck’s Depression [31] and Anxiety [32] Inventories, 3) no indications of abnormal sleep, as assessed by the Pittsburgh Sleep Quality Index [33]; 4) not considered extreme morning or evening types, as quantified with the Horne & Ostberg chronotype questionnaire [34]; and, 5) free of psychoactive or sleep-affecting medications.

Twenty-four older participants initiated the study protocol. Participation was terminated if sleep duration during the habituation nap was insufficient (less than 10 minutes; N = 3) or if the Montreal Cognitive Assessment [35] revealed signs of cognitive impairment (i.e., score below 26 ; N = 2). An additional participant was excluded due to experimental error and one individual did not complete the entire protocol. In total, 17 participants completed the study and were included in data analyses (see participant characteristics in Table 1). Note that three additional participants were excluded from *only* the analyses with detected SWs (i.e., sleep event detection and the SW-locked phase amplitude coupling described below) due to no detected SWs during epochs of interest (see below for details).

**Table 1.** Participant characteristics

|  |  |
| --- | --- |
| N | 17 (8 females) |
| Age (yrs) | 62.7 ranging from 50 to 74 |
| Edinburgh Handedness | 95.2 [88.7 - 100] |
| Epworth Sleepiness Scale | 5.4 [3.7 - 7] |
| Beck Depression Scale | 1.7 [0.3 - 3] |
| Beck Anxiety Scale | 1.6 [0.3 - 3.1] |
| PSQI | 3.4 [2.4 - 4.4] |
| Chronoscore (CRQ) | 57.9 [53.8 - 62] |
| MOCA | 28.8 [28.2 - 29.4] |

*Notes.* Values are means [lower and upper limit of the 95% Confidence Interval (CI)]. PSQI = Pittsburgh Sleep Quality Index; CRQ = Circadian Rhythm Questionnaire; MOCA = Montreal Cognitive Assessment.

### 2.2. General design

This study employed a within-participant design (Figure 1). First, to increase the probability of falling asleep during the experimental nap, all participants completed a 90-minute habituation nap monitored with polysomnography (PSG, see below for details) in the early afternoon. Approximately one week later, participants returned to the laboratory to complete the experimental protocol. Participants were instructed to maintain a regular sleep/wake schedule for the 3 days leading up to this experimental session (see Table 2 for results based on averaged data from the objective actigraph and subjective sleep diary data acquired over this interval). On the first experimental day (day 1), two motor sequences were learned simultaneously in the pre-nap session. A different sound was associated to each of these two sequences during task performance. During the subsequent nap episode, one of these two sounds was presented. This sound is referred to as the *associated* sound that was linked to the *reactivated* sequence. The sequence that was learned during the pre-nap session but not reactivated during the subsequent nap interval served as the control (*non-reactivated*) condition at the behavioral level. At the electrophysiological level, the control condition consisted of a new sound played during the nap. This sound was thus not linked to any learned material (*unassociated* sound). The experimental nap was monitored with PSG. Online monitoring of the sleep data was performed by an experimenter, affording the opportunity to send auditory stimulations during NREM2-3 stages (see below for details). Thirty minutes after the end of the nap, performance on the reactivated and non-reactivated sequences was tested (i.e., post-nap retest). The following morning (day 2), after a night of sleep spent at home (monitored with actigraphy and sleep diary; see Table 2), performance on the two sequences was again assessed (i.e., post-night retest). Objective (Psychomotor Vigilance Task [36]) and subjective (Stanford Sleepiness Scale [37]) measures of vigilance were assessed at the beginning of each testing session (results presented in Table 2). General motor execution was also tested with a random variant of the motor task (described below) at the beginning of the pre-nap session and at the end of the post-night session.

**Table 2.**
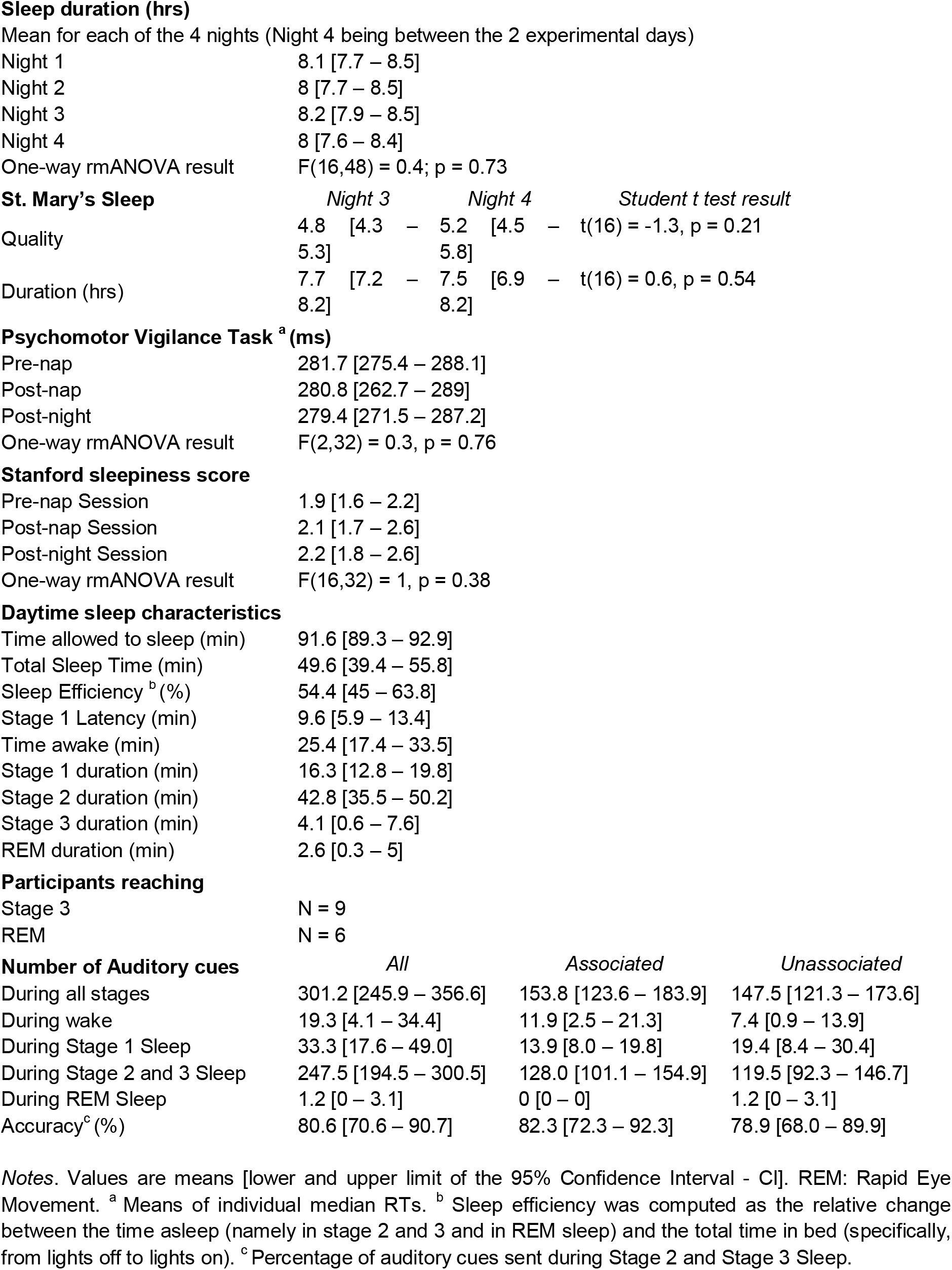
Sleep and vigilance scores (N=17)

**Figure 1:**
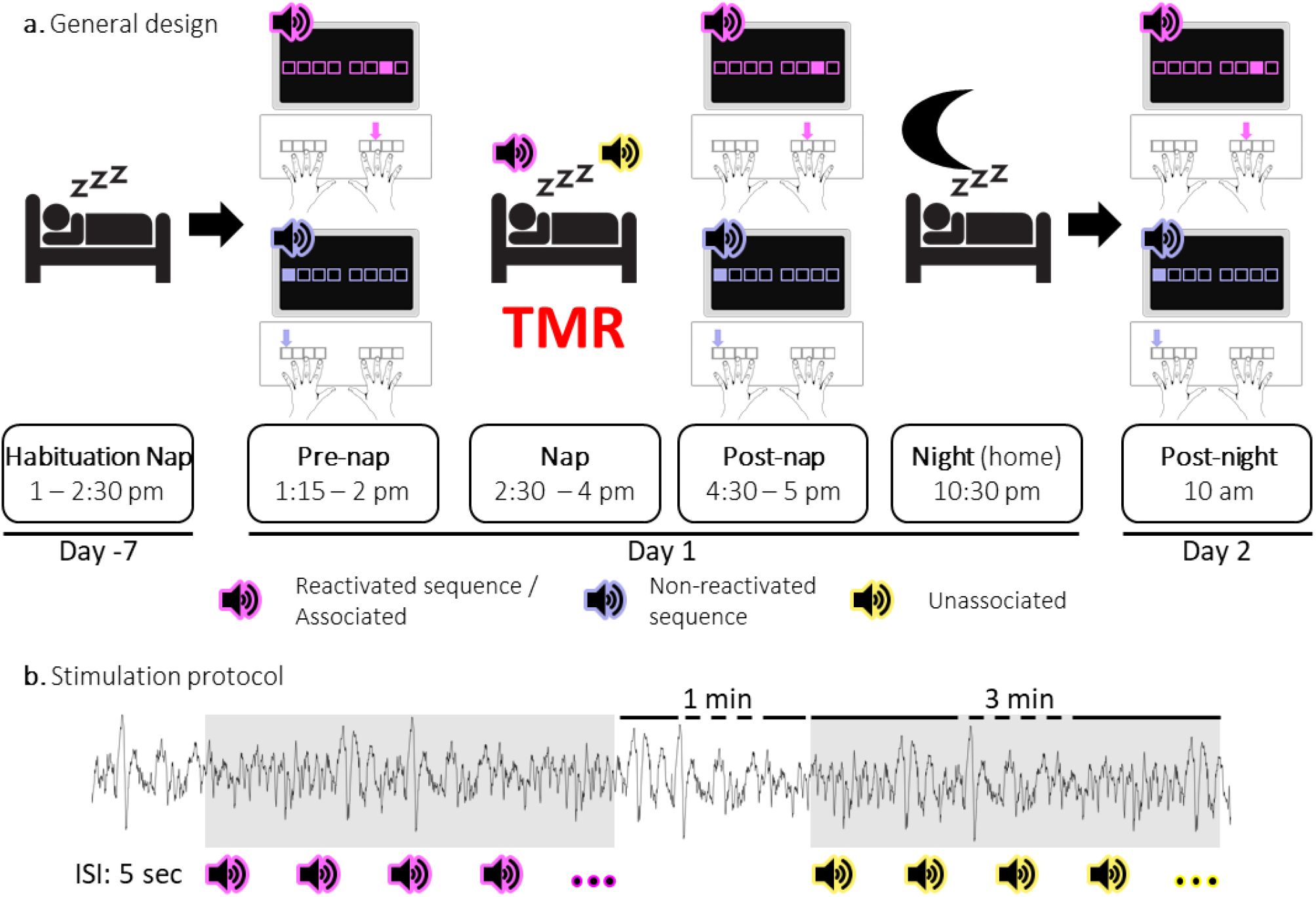
Experimental protocol. (Figure and caption were adapted from [27] under the Creative Commons Attribution (CC BY) license. **a. General design**. Following a habituation nap that was completed approximately one week prior to the experiment, participants underwent a pre-nap motor task session, a 90-minute nap episode monitored with polysomnography during which targeted memory reactivation (TMR) was applied and a post-nap retest session. Participants returned to the lab the following morning to complete an overnight retest (post-night). The times provided represent the schedule for an exemplar participant. During the motor task, two movement sequences were learned simultaneously and were cued by two different auditory tones. For each movement sequence, the respective auditory tone was presented prior to each sequence execution (i.e., one tone per sequence). One of these specific sounds was replayed during the subsequent sleep episode (**Reactivated**) and the other one was not (**Non-reactivated**). During the NREM 2-3 stages of the post-learning nap, two different sounds were presented. One was the sound associated (**Associated**) to one of the previously learned sequences, i.e., to the reactivated sequence, and one was novel, i.e., not associated to any learned material (**Unassociated**). **b. Stimulation protocol**. Stimuli were presented during three-minute stimulation intervals of each cue type alternating with a silent 1-minute period (rest intervals). The inter-stimulus interval (ISI) was of 5 sec. The stimulation was manually started when participants reached NREM sleep and stopped when participants entered REM sleep, NREM1 or wakefulness.

With this design, the effect of TMR on consolidation was assessed at the behavioral level by comparing the changes in performance between the reactivated and non-reactivated sequences and at the electrophysiological level by comparing the neurophysiological responses to the associated (i.e., reactivated) and unassociated auditory cues.

### 2.3. Stimuli and tasks

#### 2.3.1. Acoustic stimulation

During the nap, sound presentation was conducted using speakers positioned at the bed of the participant. During motor task sessions, the speakers were positioned on the desk, facing the participants on both sides of the computer. Three different 100-ms sounds (see [27] for details on sound characteristics) were pseudo-randomly assigned to the three conditions (reactivated/associated, non-reactivated, and unassociated) for each participant to ensure that the associations between sounds and conditions were evenly distributed across participants. At the start of the experiment, the individualized auditory detection threshold of each sound was determined using a transformed 1-down 1-up staircase procedure [38, 39] with the speakers positioned in the nap configuration and the participant lying down in the bed. The sound pressure level was set to 1000% of the individual auditory threshold during motor task performance and at 140% of the threshold during the nap, thus limiting the risk of awakening [40].

#### 2.3.2. Motor Task

Motor learning and memory consolidation processes were probed using a bimanual serial reaction time task (SRTT) [41]. In the present study, the SRTT consisted of eight squares presented horizontally on the screen meridian. Each square corresponded to one of the 8 fingers (no thumbs) associated to one of the eight keys on a specialized keyboard. During blocks of task practice, participants were instructed to press the key matching the location of a green filled square that appeared on the screen as fast and as accurately as possible. The next square changed to green with no response-to-stimulus interval. Each practice block concluded after 64 key presses; 15-sec rest blocks followed, indicated by the outline of the squares changing from green to red.

The order of the key presses either followed a pseudo-random or a sequential pattern, the former assessing general motor execution and the latter, motor sequence learning. For the sequential SRTT, two different eight-element sequences were learned simultaneously. Specifically, each practice block consisted of four repetitions of a specific sequence (e.g., sequence A: 1 6 3 5 4 8 2 7, where 1 through 8 correspond to the left pinky to the right pinky in left-to-right progression) and 4 repetitions of the other sequence (e.g., sequence B: 7 2 6 4 5 1 8 3). Repetitions of the same sequence were separated by 1 sec-intervals while the two different sequences within the same block were separated by 2 seconds. The order of the two sequences was randomized within each block of practice. A different sound was associated to each motor sequence and was played before the first key press of the sequence to be performed. The sequence-condition (conditions reactivated and non-reactivated; sequences A and B) as well as the sound-sequence associations (sounds 1, 2 and 3; sequence A, sequence B, and control sound presented during nap) were randomized and participants were pseudo-randomly assigned to one of these combinations. For the random SRTT, the order of the eight keys was shuffled for each eight-element repetition; thus, the number of each key press was constant across all random and sequential blocks.

During the pre-nap session, participants completed 4 blocks of the random SRTT followed by the sequential SRTT, consisting of 16 blocks of training and 4 blocks of post-training test. The post-training test assessed the end of training performance and took place after a 5-min break allowing the dissipation of physical and mental fatigue [42]. Only 4 blocks of the sequential SRTT were completed during the post-nap session, avoiding extensive task practice before the final overnight retest. Last, the post-night session consisted of 16 blocks of the sequential SRTT followed by 4 blocks of the random SRTT.

A generation task was performed between the training and post-training test blocks as well as after the post-night session. The goal of this task was to assess sequence knowledge of the learned sequences as well as the strength of the association between auditory cues and sequences. Participants were instructed to self-generate the motor sequence corresponding to the presented auditory cue. Specifically, a cue was presented and the participant attempted one sequence and this process was repeated 4 times per cue-sequence pairing. The order of the starting sequence was randomized. During the generation task, participants were instructed to focus on accuracy and not be concerned with the speed of their responses. The percentage of correct key presses (i.e., key pressed in its correct ordinal position) was computed per attempt and per sequence. Generation accuracy was computed by averaging across attempts separately for each time point (pre-nap and post-night sessions) and sequence.

### 2.4. Polysomnography and Targeted Memory Reactivation protocol

The habituation and experimental naps were monitored with a digital sleep recorder (V-Amp, Brain Products, Gilching, Germany; bandwidth: DC to Nyquist frequency) and digitized at a sampling rate of 1000 Hz. Standard electroencephalographic (EEG) recordings were made from Fz, C3, Cz, C4, Pz, and Oz according to the international 10-20 system. A2 was used as the recording reference and A1 as a supplemental individual EEG channel. The recording ground consisted of an electrode placed on the middle of the forehead. Bipolar vertical and horizontal eye movements (electrooculogram: EOG) were recorded from electrodes placed above and below the right eye and on the outer canthus of both eyes, respectively. Bipolar submental electromyogram (EMG) recordings were made from the chin. Electrical noise was filtered using a 50=□Hz notch. Impedance was kept below 5kΩ for all electrodes. During the habituation nap, Fz, Pz, Oz, and the vertical EOG were omitted. NREM2-3 sleep stages were visually detected during the experimental nap by a researcher monitoring the PSG recordings (guidelines from the American Academy of Sleep Medicine [43]). Auditory cues were then sent when NREM2-3 sleep stages were reached. Each type of auditory cue (associated or unassociated) was presented every 5 seconds during a 3-minute-long stimulation interval (Figure 1b). Stimulation intervals of each cue type were separated by 1-minute silent periods (rest intervals). Whenever the experimenter detected REM sleep, NREM1 or wakefulness, the stimulation was stopped manually and resumed when the participant reached NREM2-3 again.

### 2.5. Analysis

The open-source software R [44, 45] was used to perform statistical tests which were considered significant for p < 0.05. P-values between 0.05 and 0.10 are highlighted in the main text but referred to as non-significant trends. Corrections for multiple comparisons was conducted with the False Discovery Rate (FDR) when necessary [46]. In the event of the violation of sphericity, Greenhouse-Geisser corrections were applied. Repeated measures ANOVAs, Student t-tests or Wilcoxon signed-rank tests (when non-normal distribution was detected using the Shapiro-Wilk test) were used to perform contrast tests and F, t and V (or W) statistics were respectively reported. Effect sizes, calculated using G*power [47], are reported. Pearson or Spearman tests were used for correlation analyses (t and S statistics are respectively reported as well as rho values when significant). Nonparametric Cluster Based Permutations (CBP) tests [48] implemented in fieldtrip toolbox [49] were used for high dimensional time and time-frequency data analyses (e.g. ERP, TF and PAC analyses). For CBP contrast analyses, *Cohen’s d* is reported while rho is reported for CBP correlations.

#### 2.5.1. Behavior

##### 2.5.1.1. Preprocessing

Speed (correct response time - RT - in ms) and accuracy (% correct responses) were used to assess m*otor performance* on both the random and sequential SRTT. Mean RT of each block was computed separately for each condition. RTs from individual correct trials were excluded from the analyses if they were greater than 3 standard deviations above or below the participant’s mean correct response time for that block (3.9% in the random SRTT data and 1.1% in the sequential SRTT data). Consistent with our previous work [27], our primary dependent variable of interest was speed (but see Figure S2 in supplementary file for sequence accuracy data).

##### 2.5.1.2. Statistical analyses

Data from the initial training session (speed and accuracy) were analyzed to determine whether the two conditions significantly differed before the TMR intervention. Two two-way rmANOVAs were performed on the sequential SRTT performance with block (1^st^ rmANOVA on the 16 blocks of the pre-nap training and 2^nd^ rmANOVA on the 4 blocks of the pre-nap test) and condition (reactivated vs. non-reactivated) as within-subject factors. We also tested potential baseline differences between sequences A and B irrespective of the reactivation condition using similar analyses (results presented in Figure S3 in supplementary file).

*Overall performance changes* examined whether improvement in movement speed was specific to the learned sequences as opposed to general improvement of motor execution. The overall performance change of the sequential SRTT was computed as the relative change between the first 4 blocks of the pre-nap training and the last 4 blocks of post-night training collapsed across reactivated and non-reactivated sequences whereas the overall performance change of the pseudo-random version of the SRTT was computed as the relative change between the 4 blocks of the pre-nap session and the 4 blocks of the post-night session.

Our primary behavioral analysis of interest tested whether sequential SRTT offline changes differed between reactivation conditions after a nap and a night of sleep. To do so and similar to our previous research [27], *post-nap offline changes in performance* on the sequential SRTT were computed as the relative change in RT between the post-training test during the pre-nap session (namely the 3 last blocks of practice, see result section for details) and the post-nap session (4 blocks of practice). *Post-night offline changes in performance* were computed as the relative change in RT between the post-training test during the pre-nap session and the first 4 blocks of practice during the post-night session. The offline changes were computed separately for the two sequence conditions (reactivated and the non-reactivated). Positive offline changes reflect performance improvements (i.e., decreased RT) from pre-nap test to post-nap or post-night retests. A rmANOVA was performed on the offline changes in performance with condition (reactivated vs. non-reactivated) and time-point (post-nap vs. post-night) as within-subject factors.

For the purposes of brain-behavior correlation analyses and consistent with our pre-registered referent paper in young adults [27], a *TMR index* was computed as the difference in offline changes in performance - averaged across time points (no interaction between the condition and time-point factors, see results for details) - between the reactivated and non-reactivated sequences. A positive TMR index reflects an advantage for the reactivated as compared to the non-reactivated sequence.

To assess possible confounding factors impacting our analyses of primary interest, we tested whether vigilance differed between our experimental sessions. Specifically, one-way rmANOVAs were performed on both the median RTs of the Psychomotor Vigilance Task and the Stanford Sleepiness Scale score with session as 3-level factor (pre-nap, post-nap, and post-night). Moreover, a one-way rmANOVA was performed on the sleep duration (quantified via actigraphy and sleep diaries) during the 3 nights before and the night between the two experimental sessions. The results presented in Table 2 indicate that vigilance did not differ among sessions and that participants followed a regular sleep schedule prior to and between the experimental sessions.

#### 2.5.2. Electroencephalography

##### 2.5.2.1. Offline sleep scoring

Offline sleep stage scoring was performed by a certified sleep technologist according to criteria defined in [50] using the software SleepWorks (version 9.1.0 Build 3042, Natus Medical Incorporated, Ontario, Canada). Data were visually scored in 30-second epochs and band pass filtered between 0.3 and 30 Hz for EOG, 0.3 and 35 Hz for EEG signals and 10 and 100 Hz for EMG. A 50 Hz notch filter was also applied (details about scored data are in Table 2).

##### 2.5.2.2. Preprocessing

Functions supplied by the fieldtrip toolbox [49] were used to preprocess the EEG data. Data were screened manually by 30-sec epoch for cleaning. Data segments contaminated with muscular activity or eye movements were excluded. Data were re-referenced with the averaged signal from A1 and A2 and then band-pass filtered between 0.1-30 Hz.

##### 2.5.2.3. Event-related analyses

Auditory-evoked potentials and oscillatory activity were computed using the fieldtrip toolbox on down-sampled data (100 Hz). Segmentation of the data into auditory cue time-locked epochs was obtained separately for the associated and unassociated cues (from one second before to three seconds after the onset of the auditory cue with a correction for onset-trigger lags). Note that 18.1 % [95% CI: 8.7 – 28.8] of the NREM2-3 stages trials were discarded during data cleaning.

*Event-related potentials (ERP)* were extracted at the channel-level. Individual ERPs were baseline corrected with a mean amplitude subtraction computed on -0.3 to -0.1 sec relative to cue onset time window. ERP data were averaged across all 6 EEG channels as our low-density EEG montage did not allow fine topographical analyses (but see Figure S4 in supplementary file for channel level analysis). In a first step, we used CBP approaches on ERP data computed across conditions (associated and unassociated) to identify specific time windows during which significant brain activity was evoked by the auditory stimulation (i.e., where ERPs were significantly different from zero). Results showed that across conditions, ERP was significantly different from zero between 0.37 and 0.53 sec at the peak and between 1.03 and 1.35 sec at the trough of the potential (peak cluster p= 0.042 (*Cohen’s d* = 0.65); trough cluster p= 0.006 (*Cohen’s d* = 0.7)). In a second step, ERP amplitude was then averaged within these specific time-windows for the two conditions separately and compared using one-tailed paired Wilcoxon signed-rank tests. The hypothesis was that the associated, as compared to unassociated, stimulation intervals would evoke larger potential amplitudes.

*Event-related oscillatory activity* was computed by extracting the Time-Frequency Representations (TFRs) of the power spectra separately for the two experimental conditions and for each channel. The power was estimated using Hanning taper/FFT approach between 5 and 30 Hz with an adaptive sliding time window of five cycles length per frequency (Δt = 5/f; 20-ms step size). To highlight the power modulation following the auditory cue onsets, baseline relative change of power correction was performed on individual TFRs (baseline from -0.3 to -0.1 sec relative to cue onset). An average of all 6 EEG channels was then computed. TFR locked to auditory cues were compared between the associated and the unassociated conditions using a CBP test from 0 to 2.5 sec relative to cue onset and between 5 to 30 Hz.

##### 2.5.2.4. Sleep-event detection

Preprocessed cleaned data were down-sampled to 500 Hz and were transferred to the python environment. *S*low waves and spindles were detected automatically in NREM2-3 sleep epochs on all the channels with algorithms implemented in the YASA open-source Python toolbox [51]. Events were defined as SWs if they met the following criteria adapted from [52]: frequency between 0.5 and 2 Hz, a duration of the negative peak between 0.3 and 1.5 sec, a duration of the positive peak between 0.1 and 1 sec, an amplitude of the negative peak between 40 and 300 µV, an amplitude of the positive peak between 10 and 200 µV and an PTP amplitude between 75 and 500 µV (see Figure S5 in supplementary file for similar results when implementing detection algorithms for age-adapted criteria [25]). Events were defined as spindles if they met the criteria adapted from [53] including a duration between 0.5 and 3 sec and a frequency between 12 and 16 Hz. For further details on the SW and spindle detection algorithms (e.g. amplitude threshold), please see [54] for the original paper and [27] for our recent research that employed the algorithms.

SWs and spindles were detected in the stimulation intervals of both associated and unassociated cues as well as during rest intervals (i.e., NREM 2-3 epochs without auditory stimulation). Three participants did not show any SWs during the associated cue stimulation (N=1), unassociated cue stimulation (N=1) or rest intervals (N=1). These three participants were thus excluded from the analyses with detected SWs (i.e., sleep event detection and the SW-locked phase amplitude coupling described below). For detected SWs, the mean PTP amplitude (µV) as well as the density (number per total time in minutes spent in stimulation or rest intervals) were extracted for each participant and condition and average across channels (but see Table S2 in supplementary file for details on SW characteristics at the channel level). For spindles, amplitude, frequency and density were extracted for each participant and condition and averaged across channels (but see Table S3 in supplementary file for details on spindle characteristics at the channel level). One-tailed paired Student t-tests were used to compare dependent variables unless normality was violated. In these instances, Wilcoxon signed rank tests were employed (SW density, spindle density and amplitude). Performing pairwise comparisons instead of one-way ANOVAs (or non-parametric equivalent) was chosen to mimic the analytical pipeline of our earlier work [27]. The hypothesis was that the associated, as compared to unassociated, stimulation intervals would show higher values.

##### 2.5.2.1 Phase-amplitude coupling

Down-sampled (500 Hz) data of the 6 EEG channels were averaged together (but see Figure S6 in supplementary file for channel level data), and then transferred to the python environment. The Event-Related Phase-Amplitude Coupling (ERPAC) method [55] implemented in the TensorPac [56] open-source Python toolbox was used to investigate the modulation of the coupling between the phase of the 0.5-2 Hz oscillatory signal and the amplitude of the signal in the 7-30 Hz frequency range in relation to either the *auditory cue onset* or *the negative peak of the SWs* (detected on the Fz electrode). The raw data were filtered between 0.5-2 Hz and 7-30 Hz. Next, the complex analytical form of each signal was obtained using the Hilbert transform. The phase of the signal from -1 to 3 sec around the auditory cue onset and from -1 to 2 sec around the negative peak of the SWs was extracted from the filtered signal within the 0.5-2 Hz slow oscillation (SO) frequency band. The amplitude of the signal was also extracted in the windows described above and between 7 and 30 Hz with 0.5 Hz step size. For each time point in the analysis window (i.e., every 2ms), the circular-linear correlation of phase and amplitude values were computed across trials. This analysis therefore tested whether trial-by-trial differences in *slow oscillation* phase explained a significant amount of the inter-trial variability in signal amplitude in the analyzed time window and therefore affords the computation of the strength of the SO/sigma coupling dynamic along the analysis window. ERPAC computed separately for the two sound conditions (cue locked and SW-locked) and for the rest intervals (SW-locked only) were compared using CBP tests.

The preferred phase (PP), reflecting whether the amplitude of the signal in a given frequency band is modulated by the phase of the signal in another band, was also computed using tensorPac [56]. These analyses were restricted to the phase of the SO and the amplitude of the signal in the sigma band after the cue onset (associated vs. unassociated) and around the negative peak of the SW (associated vs. unassociated vs. rest). The amplitude was binned according to 72 phase slices, resulting in in 5-deg bins. The phase bin at which the amplitude is maximum is therefore defined as the preferred phase. The circular statistical analyses of the PP were performed using the CircStat toolbox [57] using Rayleigh test for non-uniformity and Watson-Williams multi-sample test for equal means.

##### 2.5.2.2 Correlational analyses

Similar to our pre-registered previous research [27], we performed correlations between the TMR index and the following EEG-derived data: (1) the difference between the densities of SWs detected during the associated and unassociated cue stimulation intervals using one-sided Spearman correlations; (2) the difference between the densities of spindles detected during the associated and unassociated cue stimulation intervals using one-sided Spearman correlation; (3) the relative change between the amplitude of the negative peak of the ERP following the associated and unassociated auditory cues using one-sided Spearman correlation; (4) the difference in auditory-locked sigma band power (0-2.5 sec relative to cue onset and from 12 to 16 Hz) between the associated and unassociated auditory cues using CBP test; and, (5) the difference between SO phase and sigma oscillation amplitude (12-16 Hz) coupling strength during the associated and unassociated stimulation intervals in relation to the cue onset and to the SW event using CBP test. For all one-sided tests, we predicted that the TMR index would be positively correlated with the EEG-derived metrics. Moreover, following the pre-registered analyses pipeline of the referent paper [27], we tested whether the TMR index was correlated to the generation accuracy during the pre-nap generation task (Pearson’s correlation).

## 3. Results

### 3.1. Behavioral data

Participants accurately performed the SRTT, as indicated by only 4.1% and 3.4% incorrect trials on the random and sequential task variants, respectively (see Figure S1 for detailed analyses). There was a strong and significant difference between the improvement in movement speed over the course of sequential SRTT practice as compared to the pseudo-random SRTT (t(16) = 8.4, p = 2.8e-7, *Cohen’s d* = 2.1, Figure 2a). This indicates that the observed performance improvements during the sequential task variant can be attributed to sequence-specific learning as opposed to general improvement of motor execution.

**Figure 2:**
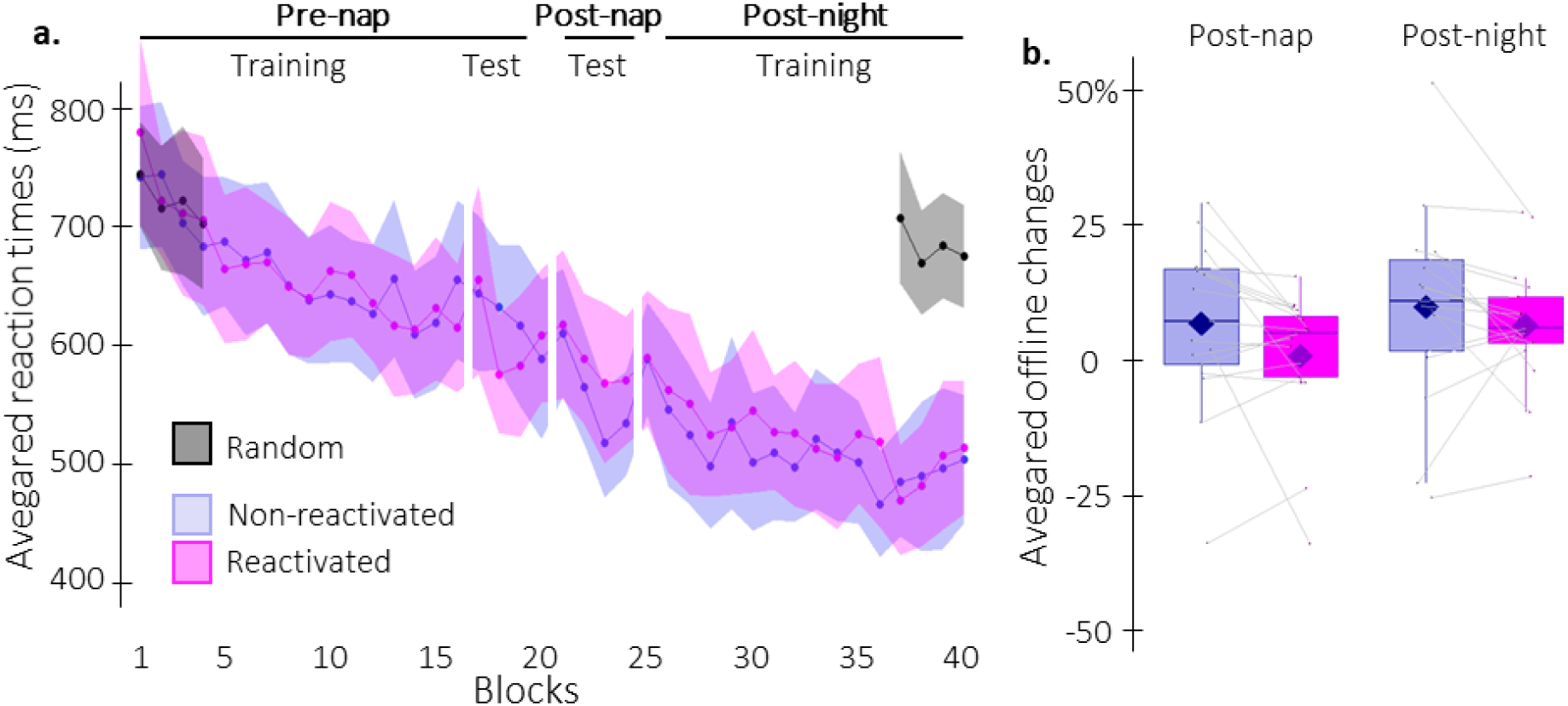
Behavioral results. **a. Performance speed** (mean reaction time in ms +/-standard error in shaded regions) across participants plotted as a function of practice blocks during the pre- and post-nap sessions for the random SRTT (black overlay) and for the reactivated (magenta) and the non-reactivated (blue) sequences. Results on performance accuracy can be found in Figure S2 in the supplementary file. **b. Offline changes in performance speed** (% change) averaged across participants for reactivated (magenta) and non-reactivated (blue) sequences and for post-nap and post-night time-points. There was no significant effect of condition (reactivated vs. non-reactivated) or condition by timepoint (post-nap vs. post-night) interaction. Box: median (horizontal bar), mean (diamond) and first(third) as lower(upper) limits; whiskers: 1.5 x interquartile range.

Analyses of the pre-nap training data indicated that participants learned the two sequence conditions (reactivated and non-reactivated sequences) to a similar extent during initial learning (16 blocks of training; main effect of block: F(15, 240) = 7.4, p = 2.42e-6, η_p_² = 0.32; main effect of condition: F(1, 16) = 5.5e-5, p = 0.99, η_p_² = 3.4e-6; block by condition interaction: F(15, 240)= 1, p = 0.41, η_p_² = 0.061; Figure 2a). Before computing offline changes in performance, we assessed whether participants reached stable and similar performance levels between conditions during the pre-nap test. Results showed that while performance reached similar levels between conditions (4 blocks; main effect of condition: F(1,16) = 1.1, p = 0.31, η_p_² = 0.063; block by condition interaction: F(3,48) = 2.1, p = 0.11, η_p_² = 0.12), asymptotic performance levels were not reached, as shown by a significant block effect (F(3,48) = 3.1, p = 0.036, η_p_² = 0.16). An identical result was observed in our reference study in young adults [27]. Similar to this previous work and to meet the performance plateau pre-requisite to compute offline changes in performance, the first block of the pre-nap test driving this effect was removed from further analyses. Performance on the remaining 3 blocks was stable, as indicated by a non-significant block effect (F(2,32) = 0.07, p = 0.93, η_p_² = 0.0045) and block by condition interaction (F(2,32)= 2.6, p = 0.087, η_p_² = 0.14). Consistent with above, the main effect of condition was not significant (F(1,16) = 1.6, p = 0.22, η_p_² = 0.092) when only including this subset of blocks. Altogether, these results indicate that a performance plateau was ultimately reached and both sequence conditions were learned similarly (Figure 2a).

For each condition separately, post-nap and post-night offline changes in performance were then computed as the relative change in speed between the three plateau blocks of the pre-nap test and the first four blocks of the post-nap and post-night retests, respectively (Figure 2b). A rmANOVA performed on offline changes in performance with time-point (post-nap vs. post-night) and condition (reactivated vs. non-reactivated) as within-subject factors highlighted a non-significant trend for a time-point main effect (F(1,16) = 3.3, p = 0.086, η_p_² = 0.17). Yet, our primary factor of interest, condition, did not reveal a significant main effect nor an interaction with time-point (condition effect: F(1,16) = 2.9, p = 0.11, η_p_² = 0.15; condition by time-point interaction: F(1,16) = 1.3, p = 0.28, η_p_² = 0.073).

These results indicate that TMR did not impact motor memory consolidation, as measured by the offline change in performance, in older adults. Offline changes were slightly, albeit non-significantly, larger at the post-night as compared to the post-nap test regardless the reactivation condition.

### 3.2. Electrophysiological data

Sleep characteristics resulting from the offline sleep scoring as well as the distribution of auditory cues across sleep stages are shown in Table 2. Results indicate that participants slept on average 49.6 minutes during the nap and that the majority of the cues (80.6%) were accurately presented in NREM sleep. The number of cues sent in the associated condition was not significantly different from the unassociated condition (t(16) = 1.2, p = 0.23). The number of analyzed trials in each condition also did not significantly differ (t(16) = 1.7, p = 0.11; mean and 95 CI for associated cues: 128.0 [101.2 - 154.9] and unassociated cues: 119.5 [92.25 - 146.7]).

#### 3.2.1. Event-related analyses

Between-condition comparisons using Wilcoxon signed-rank test showed that neither the first nor the second peak of the auditory-evoked potential had significantly different amplitude (V=53, p = 0.28, r = 0.37; V = 51, p = 0.24, r = 0.32, respectively) following associated as compared to unassociated cues. Further, CBP tests on the auditory evoked oscillatory activity did not highlight any significant clusters between the two auditory cues.

#### 3.2.2. Sleep event detection

Slow waves (SWs) and spindles were detected automatically on all EEG channels in all NREM2-3 sleep epochs. The detection tool identified on average 106.6 [95% CI: 62.1 – 151.2] slow waves and 92.7 [95% CI: 53.7 – 131.6] spindles averaged across channels during the nap episode (see Table S1 in supplementary file for the number of events detected on each channel and each condition).

Concerning the detected SWs, the peak-to-peak (PTP, Figure 3a) amplitude was not different between the three conditions (associated vs. unassociated: t(13) = -0.2, p = 0.58, Cohen’s *d* = 0.056; associated vs. rest: t(13) = 0.6, p = 0.55, Cohen’s *d* = 0.16; unassociated vs. rest: t(13) = 0.9, p = 0.37, Cohen’s *d* = 0.24). However, the density of the SWs (Figure 3b) during the associated condition was greater as compared to rest (V =96, p = 0.004 (0.012 FDR-corrected), r = 0.80). The density of the SWs during unassociated stimulation did not differ from associated stimulation or rest intervals (unassociated vs. associated: V = 73, p = 0.11 (0.16 FDR Corrected), r = 0.39; unassociated vs. rest: V = 73, p = 0.22 (0.22 FDR Corrected), r = 0.32). See Figure S5 in the supplementary file for results were when using age-adapted criteria for SW detection [25].

**Figure 3:**
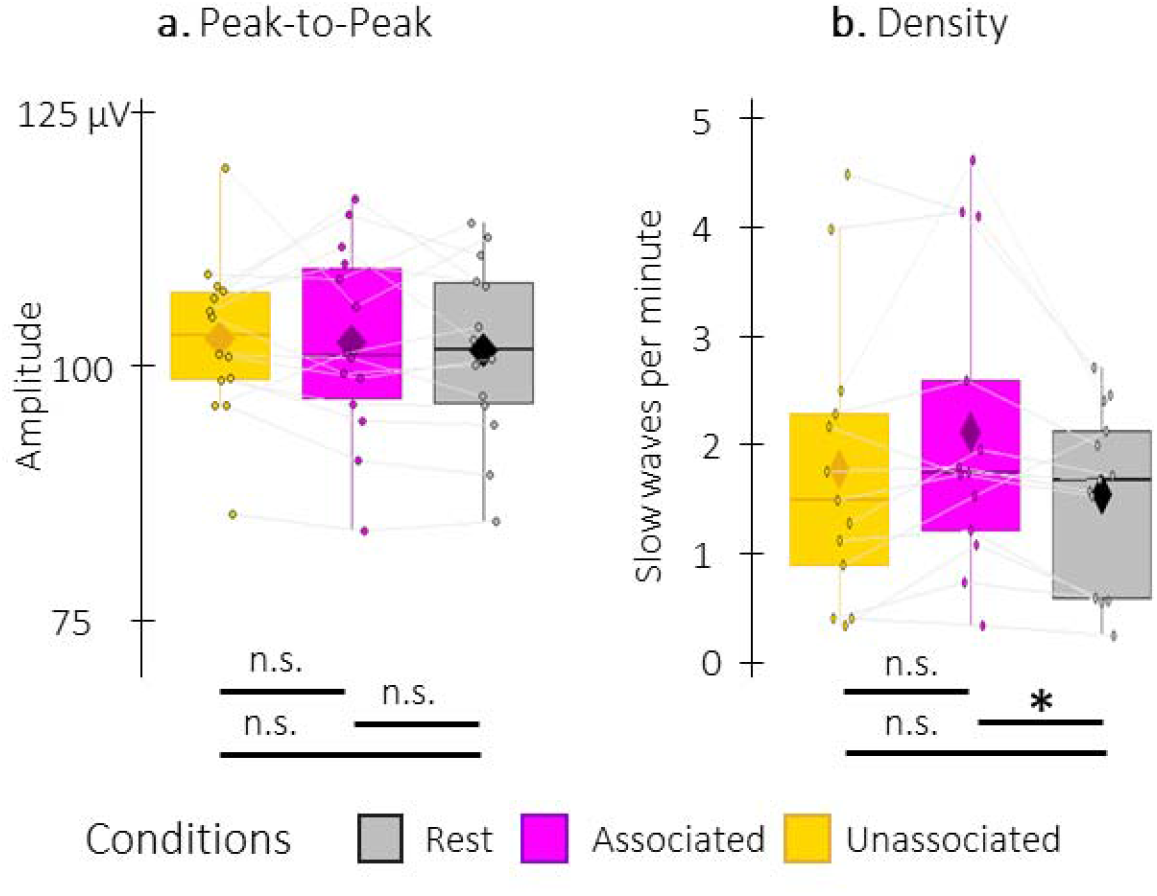
Detected Slow Waves (SWs). **a**. Peak-to-peak SW amplitude (µV) did not differ between conditions. **b**. SW density (number of SWs per total time in minute spent in stimulation or rest intervals) was higher during associated stimulation as compared to rest intervals. Box: median (horizontal bar), mean (diamond) and first(third) as lower(upper) limits; whiskers: 1.5 x interquartile range; *: p < 0.05; n.s.: not significant. For details on SW characteristics at the channel level, please refer to Table S2 in supplementary file.

Sleep spindle density, amplitude and frequency did not differ between associated and unassociated stimulation intervals (density: V = 76, p = 0.52 (0.52 FDR-corrected), r = 0.24; amplitude: V = 83, p = 0.39, r = 0.27; frequency: t(16) = 0.3, p = 0.38, Cohen’s *d* = 0.072, Figure 4). Comparisons with the rest condition showed that sleep spindle frequency did not differ between the two stimulation conditions and the rest condition (associated vs. rest: t(16) = -1.6, p = 0.13, Cohen’s *d* = 0.39; unassociated vs. rest: t(16) = -2.0, p = 0.067, Cohen’s *d* = 0.48, Figure 4b) nor did the sleep spindle amplitude (associated vs. rest: V = 102, p = 0.24, r = 0.33; unassociated vs. rest: V = 102, p = 0.24, r = 0.24, Figure 4c). However, the density of the spindles (Figure 4a) during the associated and the unassociated conditions was greater as compared to rest (V = 13, p = 0.0013 (0.0020 FDR-corrected), r = 0.70 and V = 12, p = 0.0011 (0.0020 FDR-corrected), r = 0.82, respectively).

**Figure 4:**
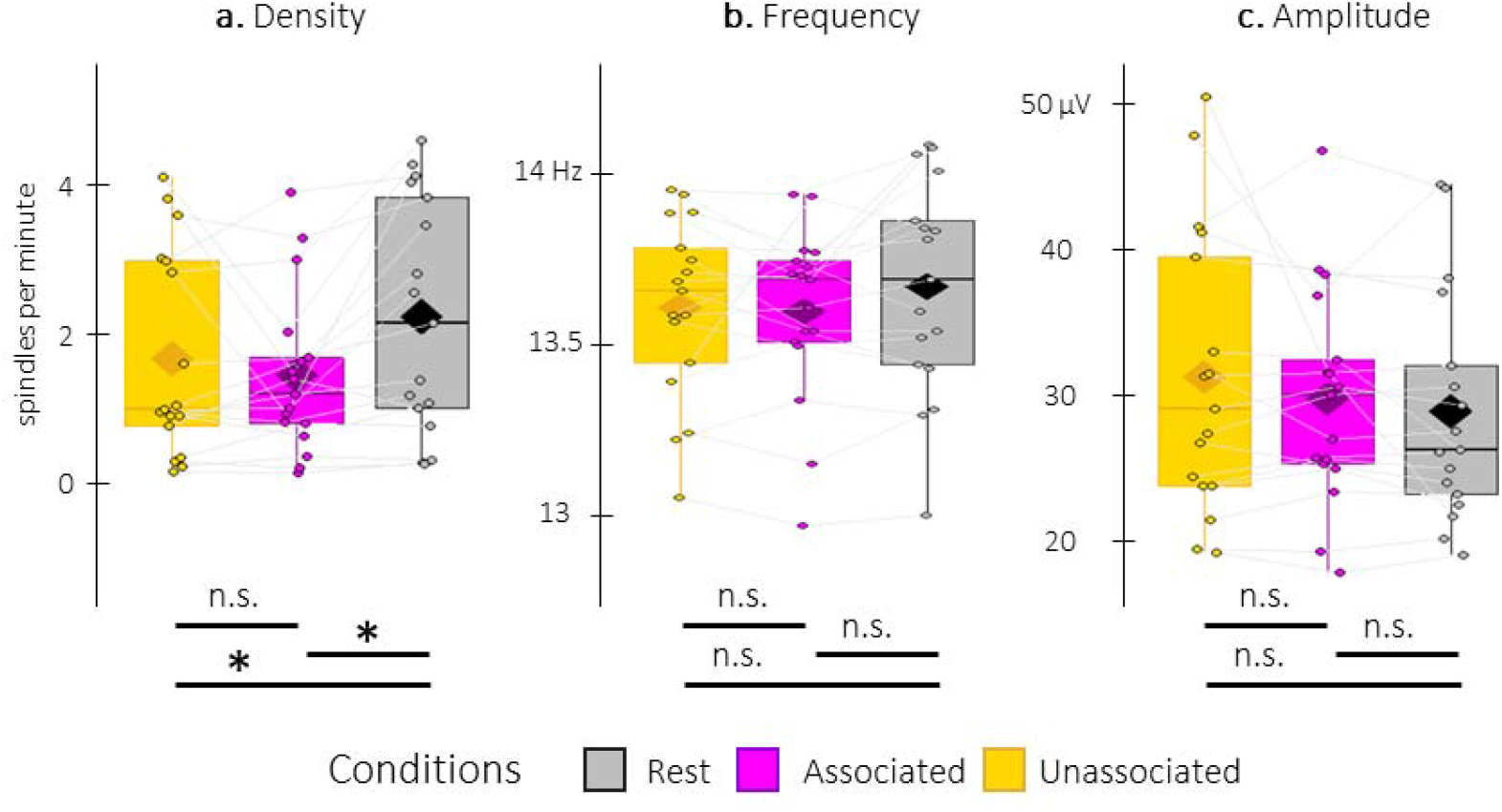
Detected spindles. **a**. Spindle density (number of spindles per total time in minute spent in stimulation or rest intervals) was lower during stimulation intervals (associated: magenta, unassociated: yellow) as compared to rest (grey) intervals. **b**. Spindle frequency (Hz) did not differ between the stimulation intervals nor as compared to rest intervals. **c**. Spindle amplitude (µV) did not differ between conditions. All spindle features were averaged across channels. Box: median (horizontal bar), mean (diamond) and first(third) as lower(upper) limits; whiskers: 1.5 x interquartile range; *: p < 0.05; n.s.: not significant. For details on spindle characteristics at the channel level, please refer to Table S3 in supplementary file.

Altogether, these results indicate that the associated stimulation resulted in an increase in SW density as compared to rest intervals but not to unassociated stimulation intervals. Furthermore, auditory stimulation – independent of whether it was associated or unassociated to previously-learned motor sequences - altered spindle density as compared to rest but not the frequency nor the amplitude of the spindles.

#### 3.2.3. Phase-amplitude coupling

We investigated whether the phase of the slow oscillations in the 0.5-2 Hz frequency band was coupled to the amplitude of sigma (12-16 Hz) oscillations following either the auditory cue or the negative peak of the detected (i.e., spontaneous) SWs. The analyses presented below focus on the comparison between conditions but see Figure S7 in supplementary file for coupling analyses performed within each stimulation condition and at rest.

The cue-locked preferred coupling phase, which represents the phase at which the maximum amplitude is observed, did not significantly differ between conditions (F(1,32)= 0.10, p = 0.75, η_p_² =0.045). This suggests that the stimulation conditions did not influence the coupling between the phase of the slow oscillations and the amplitude of sigma oscillations at the auditory cue (Figure S7 in supplementary file). Event-related phase-amplitude coupling (ERPAC) analyses performed across channels on the 7-30 Hz frequency range showed that the ERPAC values locked to the *auditory cues* did not differ between the two stimulation conditions (CBP; cluster p-values > 0.08).

The preferred phases around the negative peak of the SW were not significantly different between conditions (associated vs. unassociated: F(1,26) = 0.009, p = 0.93, η_p_² = 0.017; associated vs. rest: F(1,26) = 0.42, p = 0.52, η_p_² = 0.11; unassociated vs. rest: F(1,26) = 0.8, p = 0.39, η_p_² = 0.15; see Figure S7 in supplementary file). Comparison of the ERPAC locked to the *negative peak of the SWs* between stimulation conditions revealed no significant cluster (Figure 5a). However, a significant cluster was observed when comparing the ERPAC during associated stimulation and rest intervals (cluster threshold = 0.05, cluster p = 0.048 (above Bonferroni correction for two-tailed comparison); Cohen’s d = 0.52; Figure 5b). This cluster was observed between 20.5 and 22 Hz and -0.43 to 0.22 sec locked to the negative peak of the SW. Moreover, three significant clusters were observed between the ERPAC during unassociated stimulation and rest intervals (cluster threshold = 0.05, all cluster ps = 0.02 (below Bonferroni correction for two-tailed comparison); Figure 5c). These clusters were observed between 9 and 12 Hz and -1 to -0.02 sec (Cohen’s *d* = 0.74), between 17.5 and 20 Hz and -0.58 and 0.09 sec (Cohen’s *d* = 0.71), and between 26 and 30 Hz and -0.49 and 0.09 sec (Cohen’s *d* = 0.66) locked to the negative peak of the SW. Altogether, these results suggest that the preferred coupling phase was not modulated by the type of auditory cue. However, the coupling between slow oscillations and beta oscillations was stronger just before the onset of the SW during the associated *and* unassociated stimulation intervals. And the coupling between slow and low sigma oscillations was also stronger in unassociated as compared to rest in the same time window.

**Figure 5:**
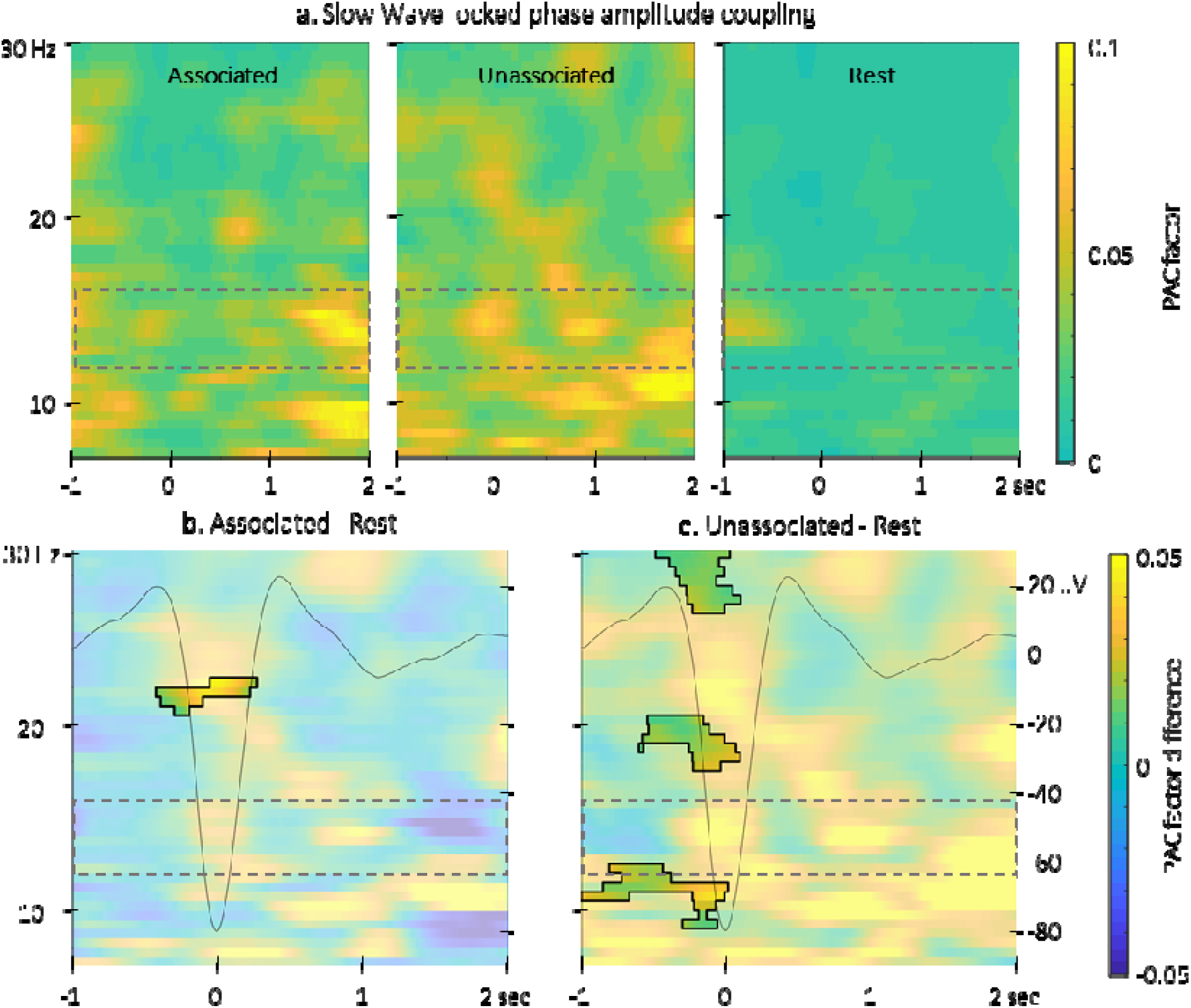
Event related phase-amplitude coupling locked to the detected slow wave negative peaks. **a**. Time-Frequency Representation (TFR) of group average coupling strength between the phase of the 0.5-2 Hz frequency band and the amplitude from -1 to 2 sec (x-axis) relative to SW negative peak and from 7 to 30 Hz (y-axis) for the three interval types. **b**. ERPAC was significantly higher around the SWs detected during the associated stimulation intervals as compared to those detected during the rest intervals in the highlighted cluster. **c**. ERPAC was significantly higher around the SWs detected during the unassociated stimulation intervals as compared to those detected during the rest intervals in the three highlighted clusters. Dashed frames indicate the sigma frequency band of interest. Superimposed on the TFR in panel b (black line): SW grand average across individuals and conditions (y-axis on right).

### 3.3. Correlational analyses

Correlation analyses between the TMR index (i.e., the difference in offline changes in performance between the reactivated and the non-reactivated sequences) and the density of either the SW or the spindles did not yield any significant results (density of spontaneous SW: S = 404, p = 0.35, r_s_ = 0.11; density of spontaneous spindles: S = 540, p = 0.09, r_s_ = 0.34). The correlational CBP analysis between the TMR index and the difference in oscillatory activity elicited by the different auditory cues did not highlight any significant clusters (all cluster ps > 0.2). Generation accuracy of the reactivated sequence during the pre-nap generation task was also not significantly correlated to the TMR index (t(15) = 0.6, p = 0.53, r = 0.16).

With respect to ERPAC-TMR index correlation analyses, no significant correlation was observed between the TMR index and the *auditory-locked* ERPAC (all cluster ps > 0.6). In contrast, CBP correlational tests performed between the TMR index and the *SW-locked* ERPAC difference (associated - unassociated) revealed a significant cluster in the 23-28 Hz frequency band and 0.47 and 1.7 sec (cluster threshold = 0.025, cluster p = 0.024, r_s_ = -0.69; Figure 6a) and a significant cluster in the 13-17-Hz frequency band and 1.14 and 2 sec (cluster threshold = 0.025, cluster p = 0.024, r_s_ = -0.63; Figure 6a) post SW-trough. The ERPAC was negatively correlated with the TMR index (cluster threshold = 0.01, cluster p = 0.001, r_s_ = -0.69; Figure 6a). For illustration purposes, we extracted the difference in ERPAC in the significant clusters. The resulting scatter plots (Figure 6b and 6c) indicate that the stronger the phase-amplitude coupling during unassociated as compared to associated stimulation intervals, the greater the TMR index. Note that, however, these results do not remain significant without the two extreme (but not defined as outlier) participants with TMR values inferior to -10.

**Figure 6.**
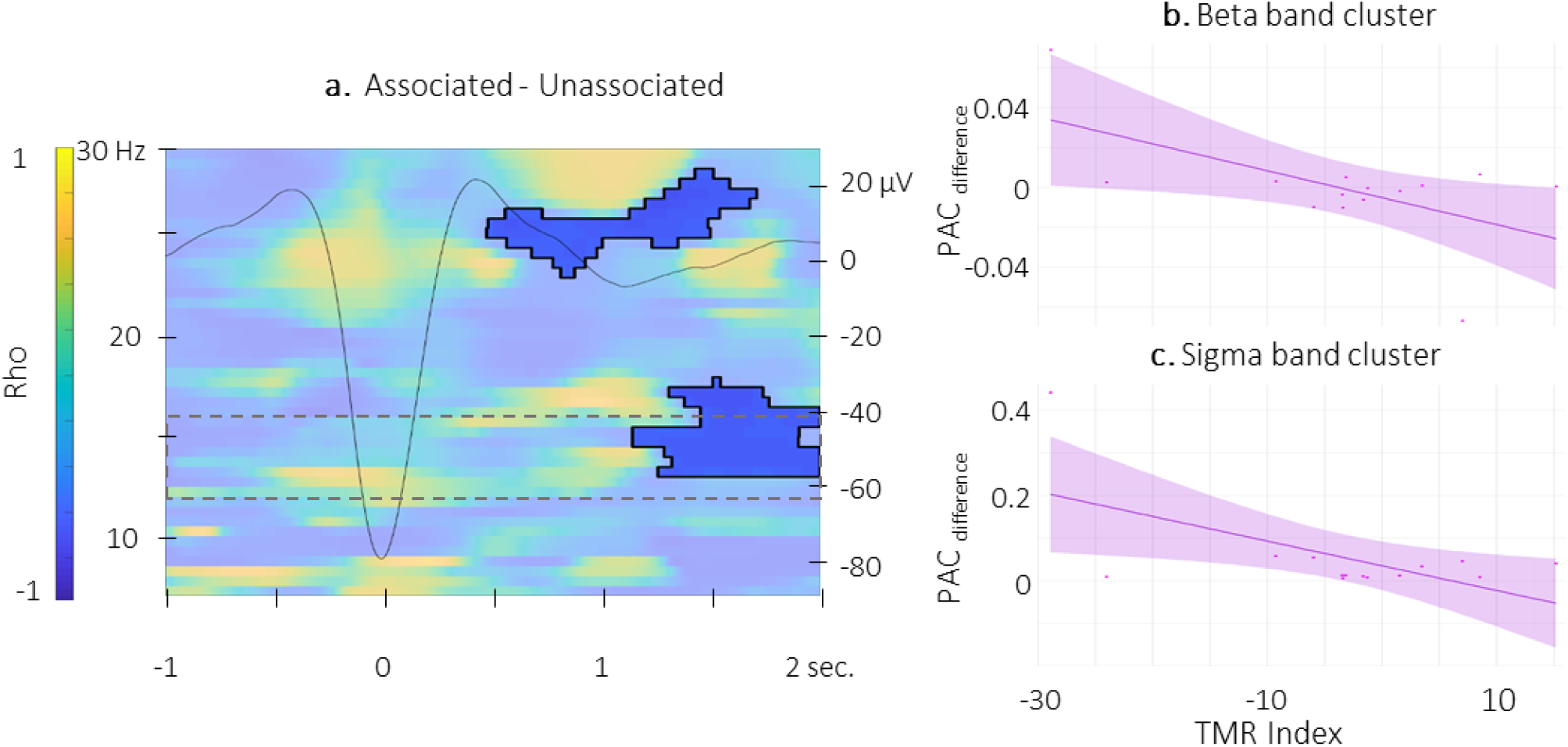
Correlation between SW-locked event related phase-amplitude coupling difference and TMR Index. **a**. Time-Frequency Representation (TFR) of the rho values resulting from the correlation between the TMR index and the difference between the SW-locked ERPAC during the associated vs. unassociated stimulation intervals (N = 14). Highlighted, the positive cluster in which the TMR index is significantly correlated with the difference in SW-locked ERPAC (cluster-based permutation test). Superimposed on the TFR (black line): SW grand average across individuals and conditions. Dashed frame highlights the sigma frequency band of interest. **b**. Depiction of the negative correlation between the SW-locked, ERPAC difference in the high beta band (0.47–1.7 s post negative peak, 23–28 Hz) and the TMR index dots represent individual datapoints). **c**. Depiction of the negative correlation between the SW-locked, ERPAC difference in the sigma band (1.14–2 s post negative peak, 13–17 Hz) and the TMR index (dots represent individual datapoints).

## 4. Discussion

This study examined the impact of auditory TMR on the behavioral and electrophysiological correlates of motor memory consolidation in older adults. Results indicated that we could not reject the null hypothesis of no TMR-induced advantage on motor performance. This lack of an effect at the behavioral level was largely paralleled by the electrophysiological results, as there were no differences between the two types of acoustic cues (i.e., associated vs. unassociated) with respect to the evoked potentials and the characteristics of the detected spindles and slow waves. Interestingly, the presentation of sounds - independent of cue type - effectively modulated the density of the spindles and the coupling between slow oscillation phase and beta band amplitude (around 20 Hz) at the descending phase of the SW. These results collectively suggest that the sleeping brain of older adults is responsive to auditory stimulation but is not sensitive to the memory content associated with different acoustic cues.

Contradictory to our predictions, results from the current study showed that sleep-related offline changes in motor performance were not modulated by TMR in older individuals. This null effect is in contrast to the TMR-induced behavioral advantage we recently demonstrated with the same protocol in young adults [27] and could potentially be attributed to a maladaptive response from the aging sleeping brain. Specifically, the sleeping brain of older adults failed to differentiate between associated and unassociated sounds, as indicated by no differences between the electrophysiological responses to the two types of acoustic cues. For example, even though significant brain activity was evoked by auditory stimulations, the evoked potentials were not significantly modulated by the type of auditory cue. Similarly, the spindle characteristics as well as the peak-to-peak amplitude of the detected SWs were equivalent in the two stimulation intervals. Additionally, although SW density was significantly higher during associated stimulation intervals as compared to rest, it was not significantly different from unassociated stimulation intervals. Interestingly, all of these electrophysiological events were modulated by the presentation of the cue associated to the memory trace in young adults [27] and thus likely reflect age-related differences.

Our electrophysiological analyses also revealed that both the associated and the unassociated cue stimulations elicited a modulation in slow/beta phase amplitude coupling as compared to rest (but did not differ between each other). Specifically, we found that PAC was higher for unassociated and associated intervals as compared to rest during the descending phase of the SW in the beta band (19 – 20.5 Hz). Interestingly, the frequency that was modulated by both sound cues in older adults was the beta band and not the sigma band which is most commonly linked to sleep-facilitated consolidation [26, 27, 58]. The role of beta in motor learning and memory consolidation processes is not without precedent, however. Previous research has demonstrated higher beta spectral power, in addition to higher sigma power and spindle activity, during sleep epochs following motor sequence learning as compared to following a control motor task [59].

Whereas the majority of our results revealed no differences in the electrophysiological responses to associated vs. unassociated stimuli during sleep in older adults, there was one exception. The difference in slow/sigma oscillation phase-amplitude coupling between the associated and unassociated stimulation intervals was negatively linked to the TMR index for two clusters. Specifically, starting at the peak of the slow oscillation, the increase in slow/high-beta (23-28 Hz) phase-amplitude coupling for unassociated (as compared to associated) sounds was related to higher TMR-induced performance enhancement. Similarly, for approximately 0.6 sec around 1.7 sec after the negative peak of the SW (1.2 sec. after the positive peak), the increase in slow/sigma phase-amplitude coupling for unassociated (as compared to associated) sounds was also related to higher TMR-induced performance enhancement. Accordingly, when assessing inter-individual differences in the electrophysiological response to the TMR intervention, there appears to be a link between associated (as compared to unassociated) phase amplitude coupling and a TMR-induced behavioral advantage. However, it is worth noting that these findings must be taken with caution, as they did not hold after removing participants with extreme, yet not outlier, TMR indices (see results).

Our data are largely consistent with the assertation that sound presentation can modulate the characteristics of specific sleep oscillations, but these neural responses are not sensitive to the memory content carried by the auditory cue, in contrast to what we previously observed in young adults. One could then speculate that the presentation of sounds during the post-learning nap boosted consolidation in older adults independently of the specific memory trace associated to the cue. The idea of a general boost of consolidation processes via sound presentation during the nap is in line with the only study, to the extent of our knowledge, that investigated TMR during sleep in older adults [60]. Their results showed that sounds boosted consolidation of all previously-acquired material (i.e., the reactivated learned material and the non-reactivated learned material) as compared to a group undergoing a nap without sounds. Thus, although our results demonstrate that the TMR intervention did not trigger enhanced consolidation of the reactivated sequence specifically, we cannot rule out the possibility that the presentation of sounds boosted sleep-related consolidation processes – across reactivated and non-reactivated sequences – in our sample of older adults. Supplemental experimental groups without any acoustic stimulation would be necessary to definitively support this explanation. Interestingly, this speculation would also be consistent with extensive research indicating that, in older adults, non-specific acoustic or non-invasive brain stimulation during a post-learning nap resulted in a memory consolidation benefit [61, 62] and altered sleep features [63].

It could be argued that the lack of effect on the performance of the reactivated sequence in older individuals is due to the lower number of stimulations delivered during the post-learning nap episode as compared to our previous research in young adults (241.6 vs. 349.5 [27]). Although feasible, we contend that this explanation is unlikely, as a recent meta-analysis reported that the beneficial effect of TMR on memory is not correlated with the number of stimulations provided during the post-learning sleep episode [12]. We reproduce this result in the current research: there was no correlation between the number of TMR stimulations and offline performance changes for the reactivated sequence (r = 0.1, p = 0.67). Another potential explanation for our pattern of behavioral results could be linked to the learning session prior to our manipulation. Specifically, it is possible that the protocol requiring participants to simultaneously learn two movement sequences mitigated initial encoding. This might have in turn compromised subsequent sleep-facilitated consolidation and consequently the impact of the TMR intervention. Such an explanation would be in line with previous research demonstrating that the initial learning process in older adults is particularly susceptible to increases in task complexity [9, 64, 65] and that the effect of post-learning sleep critically depends on performance levels achieved at the end of initial training [8, 66, 67, 68]. To further explore this possibility, we compared motor sequence learning data prior the TMR-manipulation from older adults in the current study to those acquired from the younger individuals who completed an identical protocol [27]. Results show that the learning magnitude of older adults was significantly lower than in young adults (see Figure S8 in supplementary file, for details), providing some indirect support for this explanation. To more conclusively examine the possibility that degradations in the initial encoding of older adults potentially diminished the impact of the TMR intervention, additional experimental groups consisting of older adults with extended training protocols would be necessary.

It is worth explicitly stating that as the sample in this current research can be considered moderate in size (n=17), it would certainly be of interest for future research to reproduce the reported results. Our primary behavioral contrast of interest - main effect of condition in the magnitude of offline gains - did produce a relatively large effect size (ηp² > 0.15)[69], yet the direction of this effect was opposite to our hypothesis and previous research in young adults. That is, the magnitude of offline gains in the non-reactivated condition was larger, albeit not significantly, than the reactivated condition. Accordingly, while the sample size can be considered a limitation of the current research, it is relatively unlikely that moderate increases in sample size would afford the acceptance of the alternative hypothesis that TMR *boosts* motor memory consolidation in healthy older adults.

In conclusion, in the present study, the null hypothesis was not rejected and thus it cannot be concluded that targeted memory reactivation in older adults specifically improved motor performance of the presumably reactivated motor memory trace. The electrophysiological analyses suggest that the responses to auditory stimulation during post-learning sleep are preserved in the aging brain whereas the ability to differentiate between relevant and irrelevant stimuli seems to be impaired. These findings collectively suggest that older adults do not benefit from specific reactivation of a motor memory trace by an associated sensory stimulus; however, it is possible that the presence of acoustic stimulation during post-learning sleep is an effective avenue to boost motor memory consolidation.

## Acknowledgments

This work was supported by the Belgian Research Foundation Flanders (FWO; G0D7918N), The Fond de Recherche en santé du Québec en sciences naturelles (RRQNT-2018-264146), Healthy Brain for Healthy Lives Discovery Grant Program from the Canada First Research Excellence Fund and internal funds from KU Leuven. GA also received support from FWO (G0B1419N, G099516N, 1524218N) and Excellence of Science (EOS, 30446199, MEMODYN, with SS). Financial support for authors JN was provided by the European Union’s Horizon 2020 research and innovation program under the Marie Skłodowska-Curie grant agreement (#887955).

## Data and code availability

All data can be found at data can be found at https://doi.org/10.5281/zenodo.7778543. The source code is available at https://github.com/judithnicolas/MotorMemory_OpenLoop_TMR_Aging

## Disclosure statement

The authors declare no potential conflicts of interest.

## Notes

https://doi.org/10.5281/zenodo.7778543

### Competing Interest Statement

The authors have declared no competing interest.

### Summary of Updates

After pre-processing error spotting, the manuscript has been entirely revised

