## Supplementary file for "Targeted memory reactivation during post-learning sleep does not enhance motor memory consolidation in older adults"

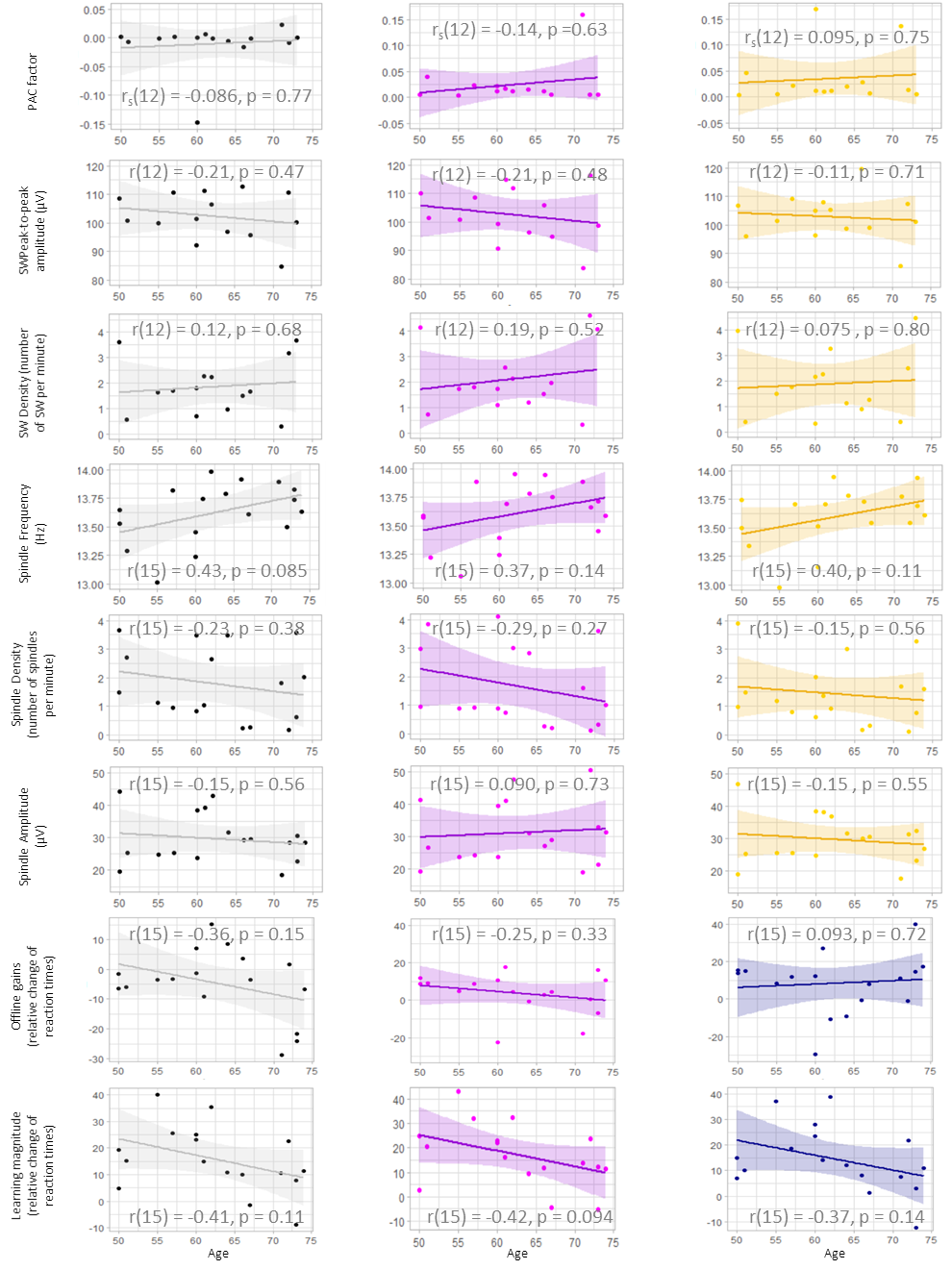
***Figure S1: Exploratory analyses assessing the correlations between the age of participants and variables of interest.*** *The different variables are displayed in separate rows while the different conditions are displayed in different columns. Rho values from Pearson (r) or Spearman (r_s_, when non-normal distribution was detected using the Shapiro-Wilk test) correlation tests are reported. For all plots, age (in years) is shown on the x-axis.* ***Grey****: collapsed across all conditions (except for PAC factor and Offline Gains where the difference between the conditions is displayed).* ***Magenta:*** *Associated stimulation (EEG variables) / reactivated sequence (behavioral variables).* ***Yellow****: unassociated stimulation (EEG variables).* ***Blue:*** *Non-reactivated sequence (behavioral variables). Note that for the correlation with PAC, values were extracted from the common significant cluster of the two conditions (namely between 17.5 and 22 Hz and from -0.58 to 0.22 sec relative to the negative peak of the slow wave). Within our sample of older adults, we did not detect any significant (i.e., p < 0.05) relationships between age and a variable of interest.*

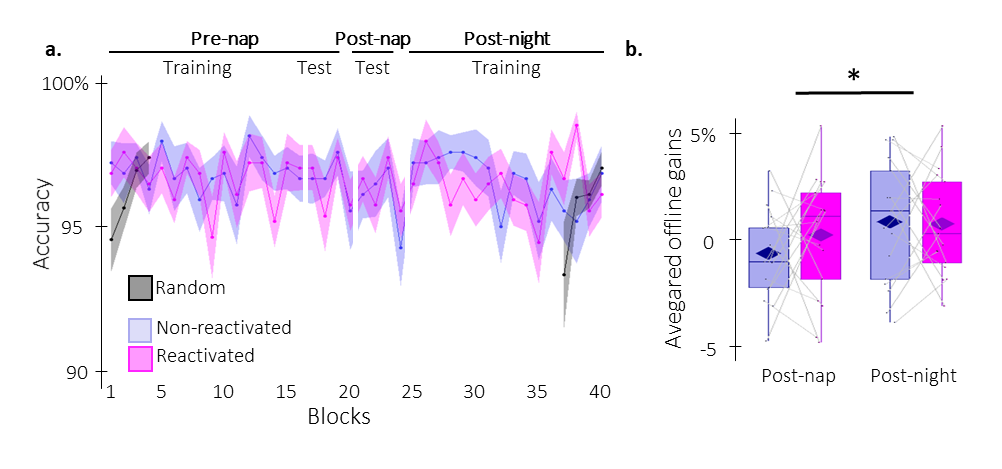

***Figure S2:*** ***Behavioral results on performance accuracy. a. Performance accuracy*** *(mean percentage of* *correct responses +/- standard error) across participants plotted as a function of practice blocks during the pre- and post-nap sessions for the reactivated (magenta) and the non-reactivated (blue) sequences. Analyses on the pre-nap training session revealed no main effects (condition: F(1,16) = 0.2, p = 0.64, η_p_² = 0.014; block: F(15,240)= 1, p = 0.39, η_p_² = 0.063) nor interaction (F(15,240)= 0.6, p = 0.87, η_p_² = 0.036). This was also the case for the pre-nap test (condition: F(1,16) =0.4, p = 0.55, η_p_² = 0.023; block: F(2,32)= 1.8, p = 0.17, η_p_² = 0.10; interaction: F(2,32)= 0.4, p = 0.68, η_p_² = 0.023).* ***b. TMR effect****. Offline changes in performance accuracy averaged across participants (box: median (horizontal bar), mean (diamond) and first(third) as lower(upper) limits; whiskers: 1.5 x interquartile range) for post-nap and post-night time-points and for reactivated (magenta) and non-reactivated (blue) sequences. Offline changes in performance accuracy were significantly higher at the post-night as compared to the post-nap time-point (F(1,16) = 6.2, p = 0.024, η_p_² = 0.28). Yet, the offline changes in performance accuracy were not different between the reactivated and the non-reactivated sequences (F(1,16) = 0.3, p = 0.6, η_p_² = 0.018). The interaction between the condition and the time-point factors was also not significant (F(1,16)= 0.8, p = 0.38, η_p_² = 0.048).*: p < 0.05.*

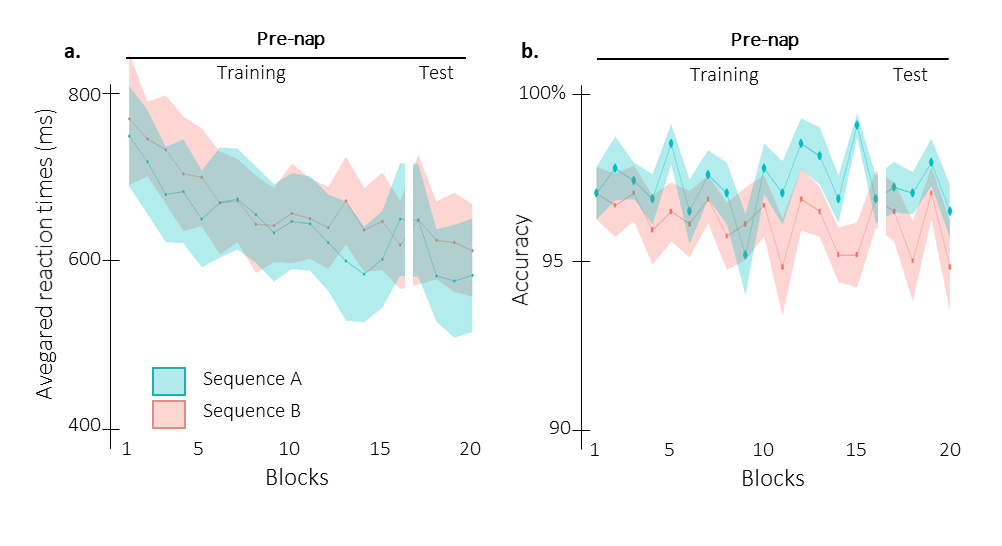
 ***Figure S3: Behavioral results per sequence during training. a. Performance speed*** *(mean reaction time in ms +/- standard error in shaded regions) across participants plotted as a function of practice blocks during the pre-nap session (training and test) for sequences A (*1 6 3 5 4 8 2 7; *green) and B (*7 2 6 4 5 1 8 3; *orange)****.*** *Analyses on the pre-nap* ***training*** *performance did not reveal a significant difference between sequences A and B (main of effect sequence: F(1,16) = 3.5, p = 0.081, η_p_² = 0.18). RT significantly decreased across blocks (main effect of block: F(15,240) = 7.4, p = 2.42e-6, η_p_² = 0.32). This decrease was not different between the two sequences (block by sequence interaction: F(15, 240)= 1.7, p = 0.11, η_p_² = 0.095). Assessment of the pre-nap* ***test*** *performance showed a significant main effect of sequence (F(1,16) = 5.6, p = 0.031, η_p_² = 0.26) but no effect of block (F(2,32) = 0.07, p = 0.89, η_p_² = 0.0045). The interaction between the block and the sequence was not significant (F(2,32)= 0.1, p = 0.89, η_p_² = 0.0071).* ***b. Performance accuracy*** *(mean percentage of* *correct responses* *+/- standard error in shaded regions) across participants plotted as a function of practice blocks during the pre-nap session (training and test) for sequences A (green) and B (orange)****.*** *The rmANOVA performed on the pre-nap* ***training*** *highlighted a main effect of sequence (F(1,16) = 11, p = 0.0044, η_p_² = 0.41), but no significant effect of the block factor nor interaction (F(15,240)= 1.1, p = 0.39, η_p_² = 0.063 and F(15,240) = 1, p = 0.42, η_p_² = 0.060 respectively). The sequence effect remained significant during the pre-nap test but no block effect or block by sequence interaction were observed (sequence: F(1,16) = 6.5, p = 0.021, η_p_² = 0.29; block: F(2,32)= 1.8, p = 0.18, η_p_² = 0.10; interaction: F(2,32)= 0.2, p = 0.82, η_p_² = 0.012). These analyses collectively revealed performance differences between the two sequences during the pre-nap session. However, as reactivation conditions were counterbalanced across sequences A and B (sequences A and B were reactivated in nine and eight participants, respectively), these effects are not considered as confounding to test our hypotheses. It is worth emphasizing that there was no significant difference between the two conditions (reactivated vs. not-reactivated) during training for both the performance speed and accuracy (see Figure 2 in the main text).*

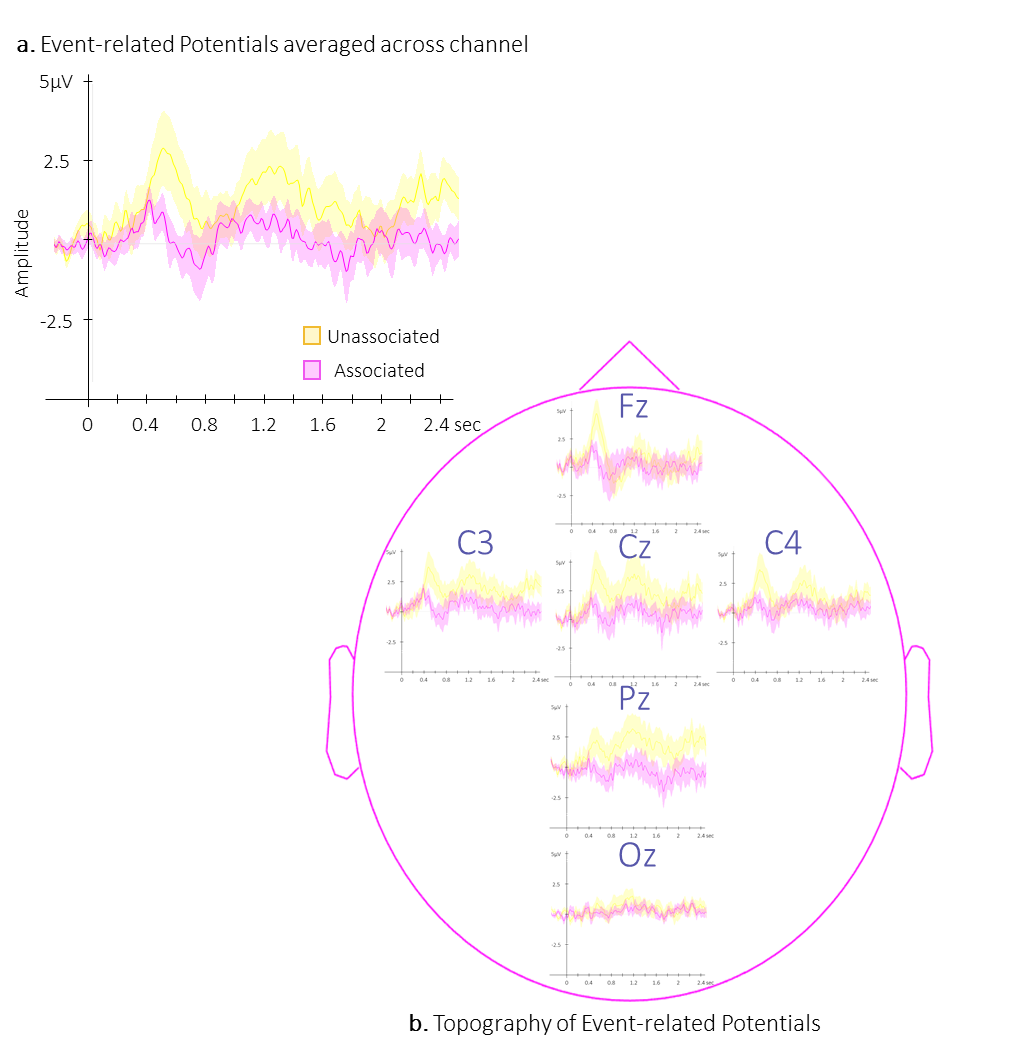

***Figure S4: Event Related Potentials*** *(ERP). Group average (+/- standard error in shaded regions) of ERP averaged across all EEG channels (****a.****) or at the channel level (****b.****) evoked by the associated (magenta) and the unassociated (yellow) auditory cues from -0.3 to 2.5 sec relative cue onset. Cluster based permutations at the channel level did not reveal any significant cluster highlighting a difference between the two conditions (associated vs. unassociated, all cluster ps > 0.27)*

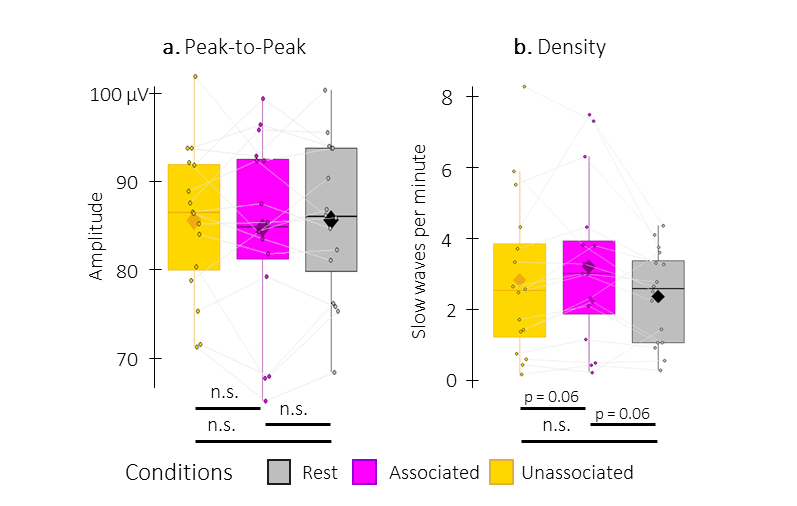

***Figure S5: Detected Slow Waves (SWs) with adapted criteria from Rosinvil and colleagues [*1*].*** *Events were defined as SWs if they met the same criteria as in main manuscript except for a peak-to-peak amplitude between 60 (instead of 75) and 500 µV and a negative peak amplitude between 32 (instead of 40) and 300 µV.* ***a.*** *Peak-to-peak SW amplitude (µV). The PTP amplitude of detected SWs were not significantly different when using the adapted criteria from [*1*] (associated vs. unassociated: t(15) = -0.5, p = 0.70, Cohen’s* d *= 0.13; associated vs. rest: t(15) = -0.8, p = 0.47, Cohen’s* d *= 0.19; unassociated vs. rest: t(15) = -0.04, p = 0.97, Cohen’s* d *= 0.01).* ***b.*** *SW density (number of SWs per minute spent in stimulation or rest intervals) was higher (trending after FDR correction) during associated stimulation as compared to rest and unassociated intervals when using the adapted criteria from [*1*]* *(associated vs. unassociated: V = 102, p = 0.042(0.06 FDR Corrected) , Cohen’s* d *= 0.35; associated vs. rest: V =112, p = 0.021 (0.06 FDR-corrected) , Cohen’s* d *= 0.64; unassociated vs. rest: V = 90, p = 0.27, Cohen’s* d *= 0.31). Note that (i) degrees of freedom are higher in this analysis as compared to the main text, as more participants could be included in the analysis (N=16) and (ii) Although not identical, these results are partially in line with those presented in Figure 3 of the main text where the density of the SWs during the associated condition was greater as compared to rest. Box: median (horizontal bar), mean (diamond) and first(third) as lower(upper) limits; whiskers: 1.5 x interquartile range; *: p < 0.05; n.s.: not significant.*

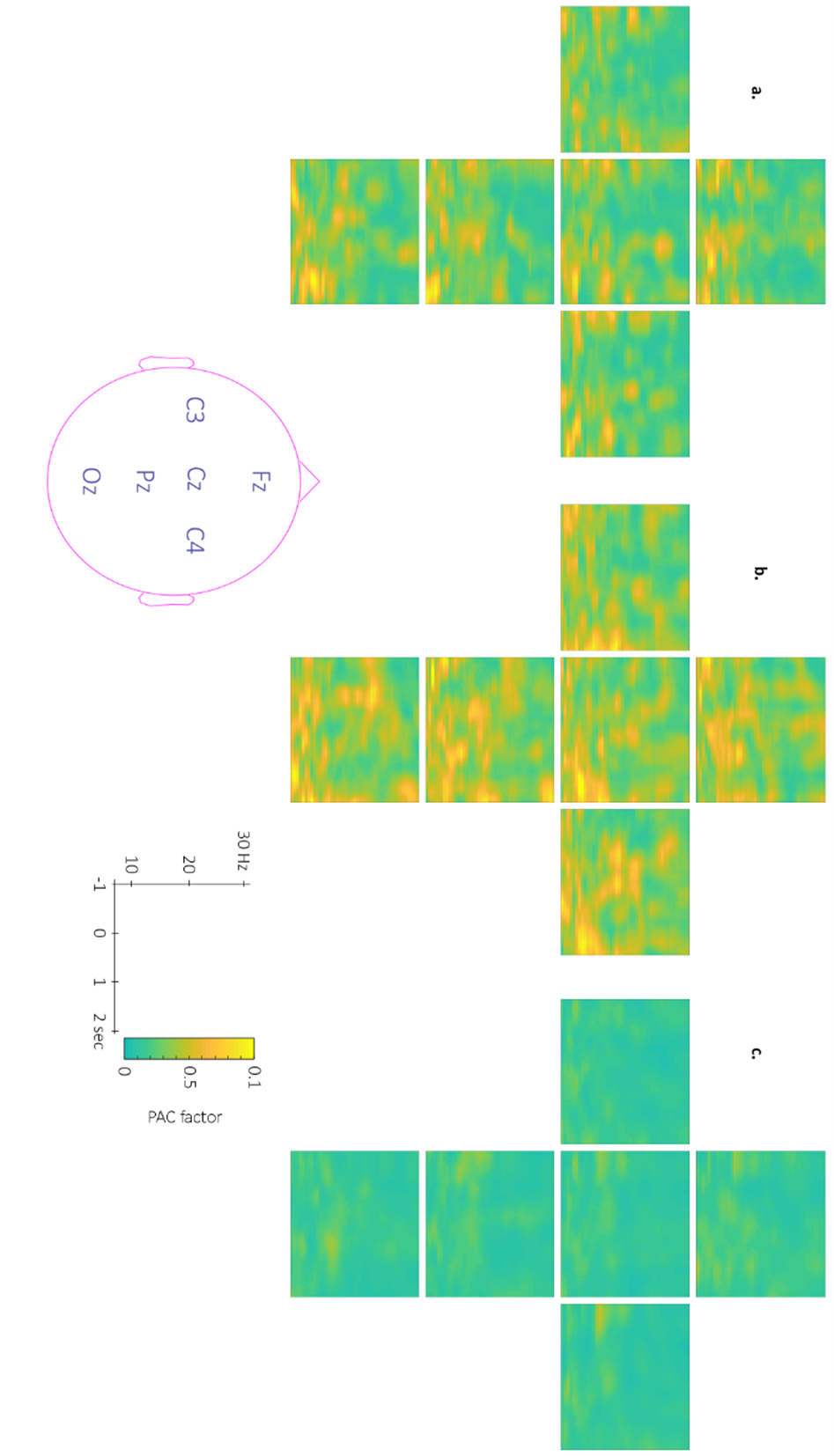

***Figure S6: Topography of the phase-amplitude coupling locked to the detected slow wave negative peaks.*** *This figure depicts the channel level data corresponding to the analyses presented in Figure 5 in the main text. SW detection was made on the Fz electrode and PAC was computed for each and every channel.* ***a-c****. Time-Frequency Representation (TFR) of group average coupling strength between the phase of the 0.5-2 Hz frequency band and the amplitude from 7 to 30 Hz (y-axis) and from -1 to 2 sec (x-axis) relative to SW negative peak for the associated stimulation (****a****.), the unassociated stimulation (****b****.) and the rest (****c****.) intervals.*

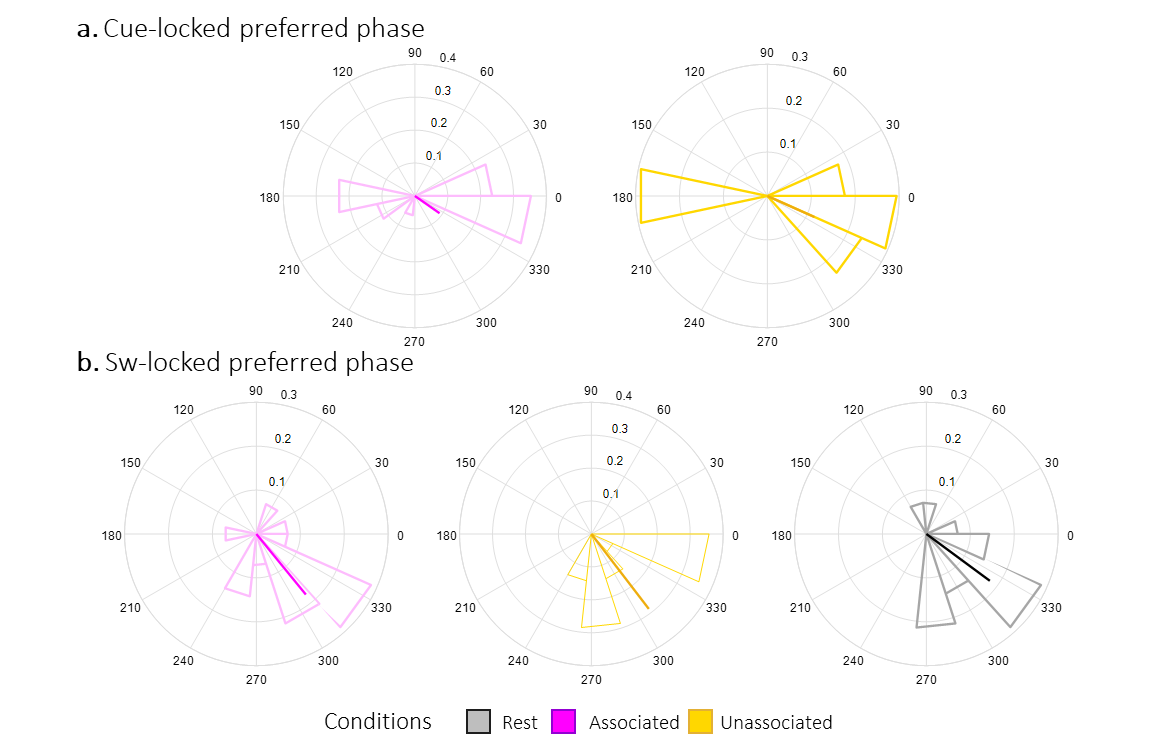

***Figure S7:*** ***Preferred Phase****.* ***a.*** *Density plot of phases in degrees (and mean direction and resultant vector length in bold) of the SO at the sigma band power peak during the 2.5 sec following associated and unassociated cues (magenta and yellow respectively).* ***b.*** *Density plot of phases of the SO at the sigma band power peak from 1 sec pre- and 2 sec post-trough of the detected SWs during associated, unassociated and rest intervals (magenta, yellow, and grey respectively). To test for the presence of a coupling between slow oscillation phases and sigma band amplitude,* *we examined whether the amplitude of the sigma oscillations peaked at a preferred phase of the slow oscillation across trials within each stimulation condition and at rest*. *To this end,* *we computed the preferred phase which is defined as the phase at which the amplitude is maximum during each periods of interest (namely each trial). Next, we tested whether the preferred phases were uniformly distributed using Rayleigh test for non-uniformity of circular data [2]; with a non-uniform distribution of the preferred phase being an indicator of coupling.* *Results failed to provide evidence that the cue-locked preferred phases across trials were distributed non-uniformly* *following associated (Rayleigh z = 1.1, p = 0.32) and unassociated cues (Rayleigh z = 2.7, p = 0.064) and thus did not demonstrate a SW-sigma coupling locked to the cue. Conversely, the phase at which the amplitude was the highest around the SW trough was distributed non-uniformly* *for SW occurring during associated stimulation (Rayleigh z = 5.4, p = 0.0029 (0.0029 FDR-corrected)), unassociated stimulation (Rayleigh z = 9.0 p = 2.29e-5, (6.87e-5 FDR-corrected)), and rest (Rayleigh z = 5.5, p = 0.0026 (0.0029 FDR-corrected)) intervals. The highest peaks of the SW-locked sigma amplitude were located on average at 312.9° [CI 95%: 296.7 - 329.1] regardless the stimulation interval. Altogether, these results indicate significant slow / sigma oscillation coupling around the SW trough but not after the auditory cue.*

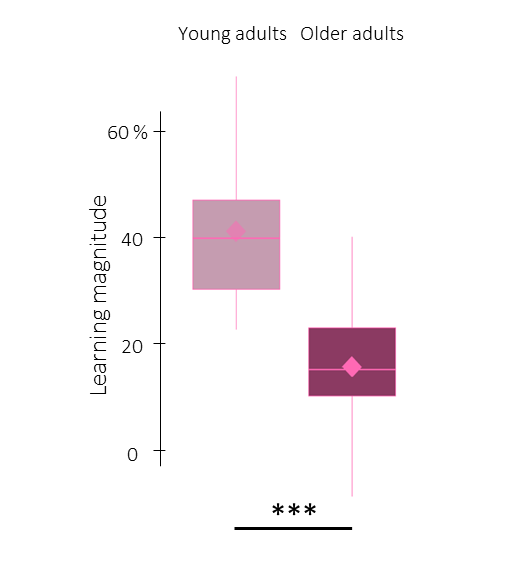
***Figure S8: Exploratory comparison of the learning magnitude between age groups.*** *To test for an effect of age on initial learning, we compared learning magnitude computed as the relative change of the sequential SRTT between the first 4 blocks of the pre-nap training and 4 blocks of pre-nap test, across sequence condition. Results show that the learning magnitude of older adults was significantly lower than the young adults one (t(35.6) = -6.4, p = 2.4e-7, Cohen’s d = 2). Box: median (horizontal bar), mean (diamond) and first(third) as lower(upper) limits; whiskers: 1.5 x interquartile range; ***: p < 0.001.*

**Table S1**. Mean number [lower and upper limit of the 95% confidence interval] of sleep events across participants detected at the single channel level per condition of intervals (either associated stimulation, unassociated stimulation or rest intervals).

|  | Slow waves | | | |
| --- | --- | --- | --- | --- |
|  | | Associated | Unassociated | Rest |
| Fz | | 53.73 [32.29 - 75.17] | 45.38 [24.8 - 65.95 | 74.31 [47.91 - 100.71) |
| Cz | | 36.5 [21.95 - 51.05] | 34.69 [15.87 - 53.51] | 46.5 [27.7 - 65.3] |
| Pz | | 21.64 [13.57 - 29.72] | 20.23 [8.36 - 32.11] | 30.57 [18.61 - 42.54] |
| Oz | | 5.5 [2.93 - 8.07] | 3.86 [1.56 - 6.15] | 4.44 [1.25 - 7.64] |
| C3 | | 25.07 [14.85 - 35.29] | 23.77 [11.92 - 35.62) | 34.73 [21.29 - 48.17) |
| C4 | | 24.14 [14.33 - 33.95) | 23.69 [12.96 - 34.42) | 32.6 [21.12 - 44.08) |
|  | | **Spindles** | | |
|  | | Associated | Unassociated | Rest |
| Fz | | 53.73 [32.29 - 75.17) | 45.38 [24.8 - 65.95) | 74.31 [47.91 - 100.71) |
| Cz | | 36.5 [21.95 - 51.05) | 34.69 [15.87 - 53.51) | 46.5 [27.7 - 65.3] |
| Pz | | 21.64 [13.57 - 29.72] | 20.23 [8.36 - 32.11] | 30.57 [18.61 - 42.54] |
| Oz | | 5.5 [2.93 - 8.07] | 3.86 [1.56 - 6.15] | 4.44 [1.25 - 7.64] |
| C3 | | 25.07 [14.85 - 35.29] | 23.77 [11.92 - 35.62] | 34.73 [21.29 - 48.17] |
| C4 | | 24.14 [14.33 - 33.95] | 23.69 [12.96 - 34.42] | 32.6 [21.12 - 44.08] |

**Table S2**. Mean amplitude and density [lower and upper limit of the 95% confidence interval] of slow waves across participants detected at the single channel level per condition of intervals (either associated stimulation, unassociated stimulation or rest intervals].

|  | Slow waves peak-to-peak amplitude (µV) | | | |
| --- | --- | --- | --- | --- |
|  | | Associated | Unassociated | Rest |
| Fz | | 116.47 [106.39 - 126.56] | 111.31 [103.97 - 118.64] | 115.87 [109.06 - 122.69] |
| Cz | | 107.24 [101.66 - 112.83] | 105.01 [98.67 - 111.34] | 103.33 [97.05 - 109.6] |
| Pz | | 100.66 [96.25 - 105.07) | 101.77 [96.09 - 107.45) | 99.75 [94.54 - 104.96) |
| Oz | | 88.55 [84.27 - 92.83) | 90.79 [85.03 - 96.54) | 90.36 [82.19 - 98.52) |
| C3 | | 102.43 [95.83 - 109.03) | 101.66 [97.03 - 106.29) | 100.46 [94.86 - 106.05) |
| C4 | | 103.22 [98.32 - 108.11) | 104.69 [99.26 - 110.12) | 100.59 [94.67 - 106.51) |
|  | | **Slow waves density (number of SW per minute)** | | |
|  | | Associated | Unassociated | Rest |
| Fz | | 3.82 [2.23 - 5.4] | 3.26 [1.94 - 4.59] | 2.79 [1.87 - 3.7] |
| Cz | | 2.62 [1.62 - 3.61] | 2.46 [1.28 - 3.63] | 1.69 [1.05 - 2.33] |
| Pz | | 1.52 [0.99 - 2.06] | 1.41 [0.62 - 2.2] | 1.08 [0.7 - 1.45] |
| Oz | | 0.36 [0.22 - 0.5] | 0.27 [0.11 - 0.43] | 0.14 [0.06 - 0.23] |
| C3 | | 1.81 [1.08 - 2.55] | 1.68 [0.95 - 2.41] | 1.27 [0.8 - 1.73] |
| C4 | | 1.72 [1.05 - 2.39] | 1.65 [0.97 - 2.34] | 1.19 [0.79 - 1.6] |

**Table S3**. Mean density, frequency, and amplitude [lower and upper limit of the 95% confidence interval] of spindles across participants detected at the single channel level per condition of intervals (either associated stimulation, unassociated stimulation or rest intervals].

|  | Spindle density (number of spindles per minute) | | | |
| --- | --- | --- | --- | --- |
|  | | Associated | Unassociated | Rest |
| Fz | | 1.25 [0.69 - 1.81] | 1.09 [0.68 - 1.51] | 1.57 [0.95 - 2.2] |
| Cz | | 1.84 [1.01 - 2.66] | 1.49 [0.86 - 2.13] | 2.22 [1.39 - 3.04] |
| Pz | | 2.6 [1.45 - 3.74] | 2.21 [1.41 - 3.02] | 3.22 [2.06 - 4.39] |
| Oz | | 1.45 [0.78 - 2.12] | 1.22 [0.64 - 1.8] | 1.91 [1.07 - 2.75] |
| C3 | | 1.73 [0.92 - 2.54] | 1.44 [0.84 - 2.03] | 2.27 [1.36 - 3.17] |
| C4 | | 1.68 [1.02 - 2.35] | 1.4 [0.79 - 2.01] | 2.42 [1.54 - 3.3] |
|  | | **Spindle Frequency (Hz)** | | |
|  | | Associated | Unassociated | Rest |
| Fz | | 13.18 [12.96 - 13.4] | 13.15 [12.99 - 13.3] | 13.32 [13.12 - 13.51] |
| Cz | | 13.68 [13.47 - 13.9] | 13.72 [13.53 - 13.91] | 13.75 [13.58 - 13.92] |
| Pz | | 13.87 [13.68 - 14.06] | 13.88 [13.66 - 14.1] | 13.93 [13.72 - 14.14] |
| Oz | | 13.8 [13.59 - 14.02] | 13.88 [13.68 - 14.09] | 13.92 [13.69 - 14.14] |
| C3 | | 13.61 [13.42 - 13.79] | 13.5 [13.33 - 13.67] | 13.61 [13.43 - 13.79] |
| C4 | | 13.48 [13.32 - 13.64] | 13.49 [13.31 - 13.67] | 13.55 [13.37 - 13.72] |
|  | | **Spindle Amplitude (µV)** | | |
|  | | Associated | Unassociated | Rest |
| Fz | | 34.16 [28.5 - 39.81] | 32.14 [27.18 - 37.11] | 32.33 [26.73 - 37.93] |
| Cz | | 34.56 [29.1 - 40.02] | 34.09 [29.31 - 38.88] | 33.38 [28.56 - 38.21] |
| Pz | | 34.23 [28.81 - 39.65] | 33.31 [28.9 - 37.72] | 31.96 [27.26 - 36.66] |
| Oz | | 19.83 [16.12 - 23.55] | 20.16 [16.41 - 23.91] | 19.8 [16.85 - 22.74] |
| C3 | | 28.91 [23.68 - 34.13] | 28.28 [23.84 - 32.71] | 27.58 [23.46 - 31.7] |
| C4 | | 28.81 [24.01 - 33.6] | 28.75 [24.81 - 32.69] | 27.76 [23.31 - 32.21] |
